# Single-cell multi-omic analysis of the vestibular schwannoma ecosystem uncovers a nerve injury-like state

**DOI:** 10.1101/2022.11.18.517051

**Authors:** Thomas F. Barrett, Bhuvic Patel, Saad M. Khan, Aldrin K.Y. Yim, Sangami Pugazenthi, Tatenda Mahlokozera, Riley D.Z. Mullins, Gregory J. Zipfel, Jacques A. Herzog, Michael R. Chicoine, Cameron C. Wick, Nedim Durakovic, Joshua W. Osbun, Matthew Shew, Alex D. Sweeney, Akash J. Patel, Craig A. Buchman, Allegra A. Petti, Sidharth V. Puram, Albert H. Kim

**Author notes:** These authors contributed equally to this work (Thomas F. Barrett, Bhuvic Patel). Correspondence can be addressed to,.

## Abstract

Vestibular schwannomas (VS) are benign tumors that lead to significant neurologic and otologic morbidity. How VS heterogeneity and the tumor microenvironment (TME) contribute to the pathogenesis of these tumors remains poorly understood. We performed scRNA-seq on 15 VS samples, with paired scATAC-seq in six samples. We identified diverse Schwann cell (SC), stromal, and immune populations in the VS TME and found that repair-like and MHC-II antigen presenting subtype SCs are associated with increased myeloid cell infiltrate, implicating a nerve injury-like process. Deconvolution analysis of RNA-expression data from 175 tumors revealed Injury-like tumors are associated with larger tumor size, and scATAC-seq identified transcription factors associated with nerve repair among SCs from Injury-like tumors. Ligand-receptor analysis and functional *in vitro* experiments suggested that SCs recruit monocytes. Our study indicates that Injury-like SCs may cause tumor growth via myeloid cell recruitment and identifies molecular pathways that may be targeted to prevent tumor progression.

## INTRODUCTION

Vestibular schwannomas (VS) are benign tumors that arise from the Schwann cells (SCs) lining the vestibulocochlear nerve and account for 8% of all primary intracranial tumors^1^. These tumors most frequently arise sporadically (> 90%) but are also associated with the autosomal dominant syndrome neurofibromatosis type 2 (NF2) and the related, but rare syndrome, schwannomatosis. Due to their anatomic location adjacent to the brainstem, both tumor growth and current treatment strategies (*i.e.,* microsurgery and/or radiation therapy) can be associated with substantial, lifelong neurologic and otologic morbidity, including hearing loss, facial palsy, disequilibrium, brainstem compression, hydrocephalus, and, in extreme cases, death^2–5^. Recent epidemiologic evidence suggests that the lifetime prevalence of VS is as high as 1 in 500 adults, largely due to incidental detection of asymptomatic tumors, which has increased with increased clinical utilization of computed tomography (CT) and magnetic resonance imaging (MRI)^6^. However, our knowledge of the molecular drivers of VS pathogenesis remains limited.

Loss-of-function mutations in the *NF2* gene are believed to be the central oncogenic event in the development of VS, but it is unknown how this genetic aberration affects downstream pathways, intercellular interactions, and expression heterogeneity *in vivo*^7–9^. First identified in patients with NF2 in the early 1990s, many studies have since sought out the pathways altered by loss of the *NF2* gene product Merlin and have demonstrated its role in a number of known oncogenic pathways *in vitro,* including Ras/Raf/MEK/ERK^10^, mTORC1/2^11^, Rac/p21-PAK/c-Jun Kinase^12^, PI3K/AKT^13^, and Wnt/β-catenin^14^. However, pre-clinical and early clinical studies of targeted inhibitors of these pathways have shown negative or, at best, modest results in limiting tumor growth^15–17^. Only bevacizumab, an anti-angiogenic agent, has been shown to limit growth in a subset of NF2 patients, but not without the risk of significant side effects^18^. Given the low burden of genomic alterations in VS, a deeper understanding of the molecular pathogenesis of VS may be advanced through detailed investigation of the transcriptional and epigenetic alterations in these tumors.

Single-cell RNA sequencing (scRNA-seq) enables characterization of the cellular compartments of tumors (*e.g.,* malignant, stromal, immune, *etc.*), as well as identification of the expression heterogeneity that exists within each of these compartments, both within and across patients^19^. More recently, single cell assay of transposase accessible chromatin sequencing (scATAC-seq) has emerged as a means for epigenetically profiling distinct cellular subpopulations, providing insights into gene regulation and determination of cell fate that complements expression data^20^. However, no study to date has described both the transcriptional and epigenomic profile of the VS TME at single cell resolution, or more broadly, utilized a multi-omic approach to study VS.

In this study, we performed scRNA-seq and scATAC-seq to characterize the expression heterogeneity and epigenetic states of cells comprising the VS TME. Within the SC compartment, we uncovered unexpected heterogeneity of SC phenotypes and found that VS-associated tumor Schwann cells (VS-SC) resemble repair-type SCs found in the setting of peripheral nerve injury. We found that a subset of tumors was enriched for repair-like cells and antigen presenting SC (“Injury-like VS”), while other tumors were characterized by low expression of these transcriptional profiles and higher expression of core markers of non-myelinating SC (“nmSC Core VS”). We also found monocytes/macrophages (herein referred to as myeloid cells) to be the most abundant immune cells in the VS TME, with their enrichment being correlated with higher fractions of repair-like and MHC II antigen presenting VS-SCs. Through deconvolution of bulk RNA-seq and expression microarray datasets, we characterized tumors with high and low myeloid cell infiltrate as Injury-like and nmSC Core and found that Injury-like tumors were associated with larger tumor size. Epigenetic analysis of VS-SCs in these distinct tumor states identified regulatory transcription factors (TFs; *e.g.*, RUNX2, FOSL1, FOSL2) that are also expressed in the setting of peripheral nerve injury. Lastly, we explored the interactions between VS-SC and myeloid cells to identify candidate targets that might disrupt these interactions.

## RESULTS

### Single cell transcriptional and epigenetic profiling identifies cellular diversity across the vestibular schwannoma tumor ecosystem

We performed scRNA-seq transcriptional profiling of 15 sporadic VS with paired scATAC-seq profiling of six tumors to capture a detailed portrait of the human VS tumor ecosystem (Figure 1A-B). After correcting for ambient RNA and removing doublets, low quality cells, and lowly expressed genes, we retained 112,728 high quality cells and 9,524 genes for downstream transcriptional analysis, and 31,578 cells with a median of 5,957 fragments per cell for downstream epigenetic analysis (Figure 1C-D, S1A).

**Figure 1.**
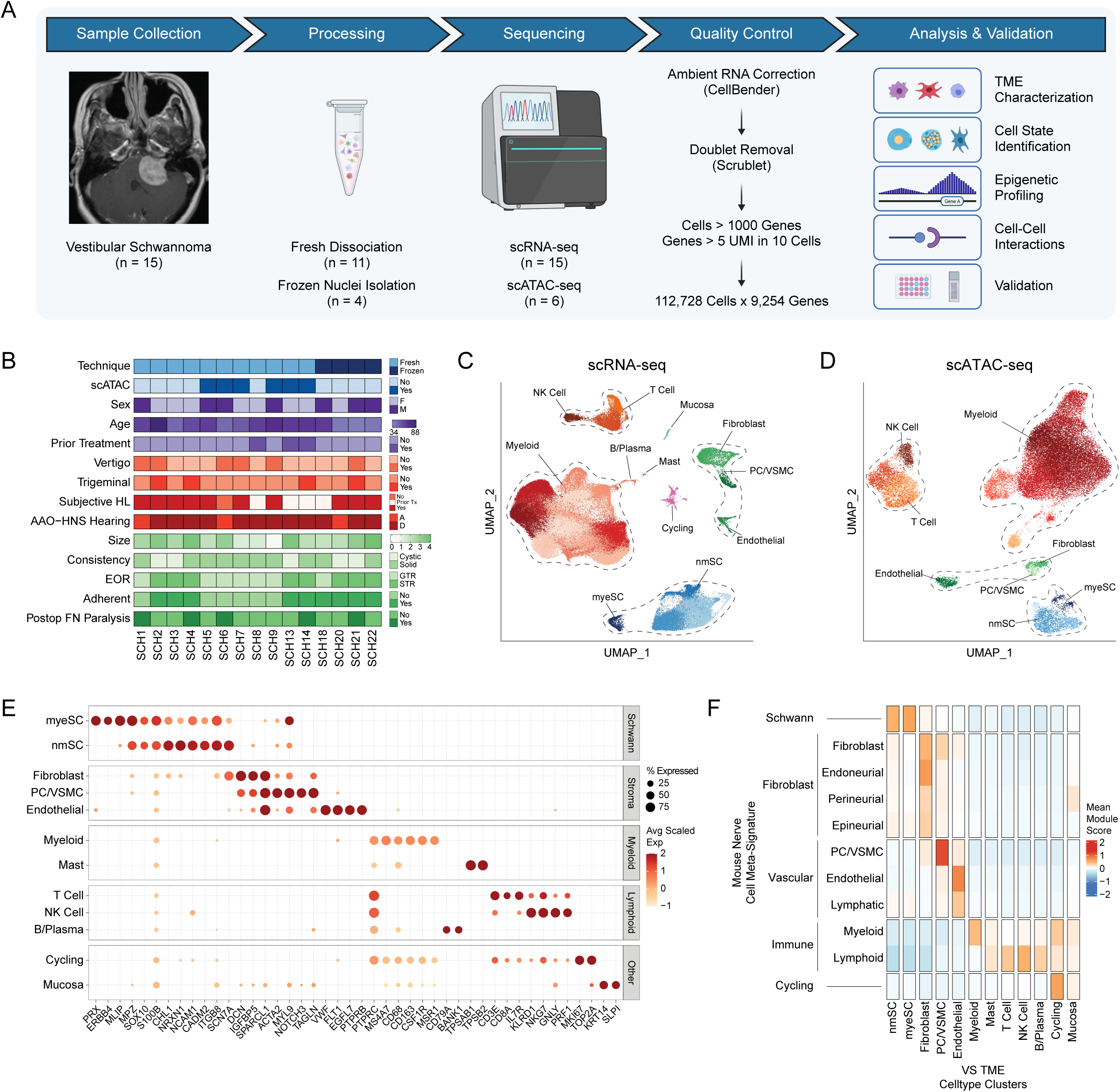
scRNA-seq and scATAC-seq atlas of vestibular schwannoma (VS). (A) Schematic of study design. (B) Clinical and demographic characteristics of tumors included in scRNA-seq and scATAC-seq datasets. AAO-HNS Hearing, American Association of Otolaryngology Head and Neck hearing score; EOR, extent of resection; FN, facial nerve. Size, greatest axial dimension in cm. (C) UMAP plot of cell types identified in the VS TME via scRNA-seq analysis. NK, natural killer cells; VSMC, vascular smooth muscle cells; nmSC, non-myelinating Schwann cells; myeSC, myelinating Schwann cells. (D) UMAP plot of cell types identified in the VS TME via scATAC-seq. (E) Dot plot of expression levels of selected marker genes (x-axis) for each VS cell subpopulation depicted in (C, y-axis). (F) Heatmap of meta-signature scores from gene signatures of previously published mouse peripheral nerve studies (see also Figure S1B).

We first assigned cell-type labels to cells within the scRNA-seq dataset using a cluster-based approach. We annotated clusters using differentially expressed genes and visualized them with Uniform Manifold Approximation and Projection (UMAP) (Figure 1C). This analysis revealed five overarching classes of cells: Schwann cells, fibroblasts, vascular (*e.g.*, pericytes and endothelial cells), immune (*e.g.*, monocytes/macrophages, T cells, NK cells, and small populations of mast cells and B cells) and cycling cells. One additional cluster was characterized by expression of epithelial markers (*KRT1, SLPI*) and was almost exclusively derived from one tumor (SCH4). These cells were likely derived from temporal bone mucosa in the surgical field that were incidentally captured during specimen collection and were therefore excluded from further analysis. Among tumor SCs, there were two distinct clusters: One characterized by typical markers of myelinating SCs (myeSC), including *PRX* and *MPZ* ^21^, and another, larger SC cluster expressing genes associated with VS and a non-myelinating SC identity (nmSC), including *S100B, SOX10*, *NRXN1, SCN7A* with lack of *PRX* expression (Figure 1E)^22^. To confirm our cell type classifications, we scored all cells in our data with gene signatures derived from published scRNA-seq peripheral nerve transcriptomic atlases^21,23–26^. We found strong concordance between our cell-type labels and both the individual prior study labels (Figure S1B) as well as the aggregated meta-signature scores for these cell-type signatures (Figure 1F).

Next, we analyzed the six samples with paired scATAC-seq data. After filtering for low quality cells and doublets (Figure S2A-C), we performed dimensionality reduction (Figure 1D) and an initial cluster-based analysis using marker genes derived from gene accessibility, as was performed with scRNA-seq data (Figure S2D). Unconstrained pairing of scRNA-seq cells with cells in the scATAC-seq atlas based on shared transcriptional and gene score profiles showed excellent overlap with the *a priori* scATAC cluster-based assignments (Figure S2E-H), suggesting that we retained all major VS TME cell-type classes in the scATAC-seq data and allowing us to reliably perform integrative downstream analysis combining transcriptional and epigenetic data on an individual cell basis.

### VS-SC adopt diverse functional states

VS typically carry a low tumor mutational burden, with the most common genetic aberrations being *NF2* loss of function mutations and loss of chromosomal arm 22q^27^. We inferred copy number alterations (CNA) of single cells using CONICSmat^28^. As expected, we observed enrichment for CNA within the SC clusters relative to other cell types (Figure 2A) consistent with these being the tumorigenic cells of VS. Notably, 22q loss was observed in 4 of our 15 samples and was almost exclusive to the nmSC cluster. Based on our clustering analysis, SCs harboring 22q loss did not significantly differ transcriptionally from cells without 22q loss, suggesting that VS-SC functional states may converge downstream of initial mutagenic events.

**Figure 2.**
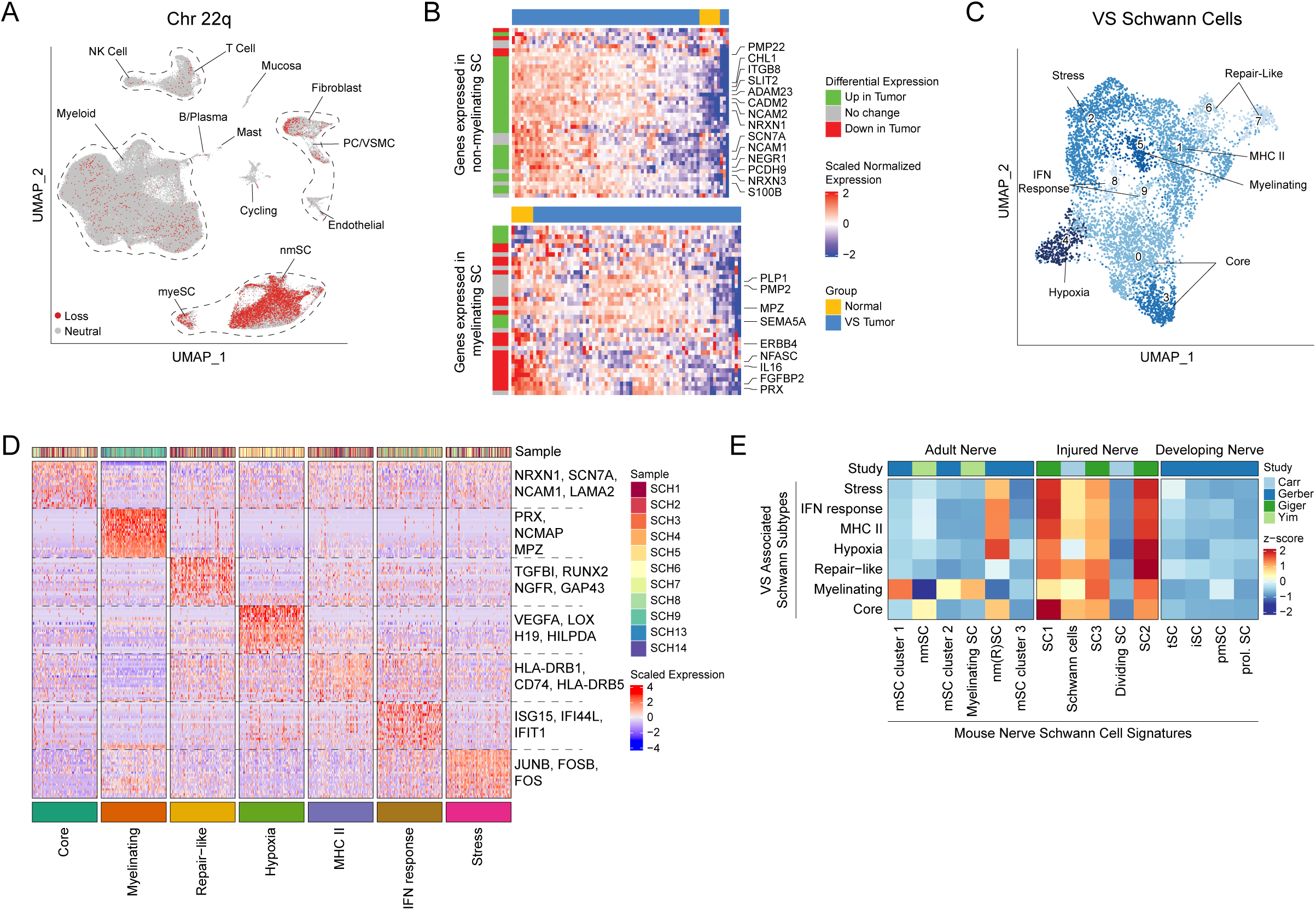
VS Schwann cells have heterogeneous transcriptional profiles. (A) UMAP plot of scRNA-seq VS data highlighting cells harboring inferred chromosome 22q (Chr 22q) loss showing enrichment in the nmSC and myeSC clusters. nmSC, nonmyelinating Schwann cells; myeSC myelinating Schwann cells. (B) Heatmaps comparing expression of top 50 differentially expressed genes (DEGs) in nmSC (top) and myeSC (bottom) to expression observed in microarray data of normal nerve and VS tumors from Gugel *et al.* (GSE141801). (C) UMAP representation of VS Schwann cells subset from the scRNA-seq data with meta-clusters labeled. (D) Heatmap of expression of DEGs from each SC meta-cluster. Two-hundred randomly sampled cells from each meta-cluster are displayed. (E) Heatmap depicting scoring of each VS Schwann cell cluster using signatures from murine adult normal nerve, adult injured nerve and developing nerve scRNA-seq atlases.

Next, we obtained publicly available RNA microarray expression datasets that compared gene expression in VS samples relative to control nerves (n = 125 tumor samples; GSE141801^29^, GSE39645^30^, and GSE108524^31^) and compared expression of the top 50 differentially expressed genes (DEGs) defining the nmSC and myeSC clusters between tumors and normal nerves in the microarray data (Figure 2B, Figure S3A). The gene signature defining VS-nmSC was markedly enriched in tumors relative to normal nerves across all 3 datasets, consistent with prior work suggesting VS-SC lose their differentiated, myelinating phenotype in favor of a less differentiated, non-myelinating phenotype^32^. Interestingly, there was mixed upregulation and downregulation of VS-myeSC associated genes in tumors relative to normal nerve controls, with a notable decrease in expression of canonical myelination markers (*e.g.*, *PRX, MLIP, NFASC, NCMAP, FGFBP2*). The mixed expression pattern of myeSC markers in tumors relative to normal nerve may represent the capture of normal bystander myeSCs or may suggest that VSs harbor a subpopulation of SCs that exist in an intermediate state before losing their myelination phenotype. Overall, this analysis served as further evidence that the VS-SC in the scRNA-seq data were indeed the tumorigenic cells of interest.

We next sought to characterize the functional states of the VS tumor SCs both *within* and *across* tumors. We selected the myeSC and nmSC clusters from the full scRNA-seq dataset and reanalyzed them by performing dimensionality reduction and batch correction, revealing ten SC subclusters, which we narrowed down to eight meta-clusters based on transcriptional similarities identified using hierarchical clustering (Figure 2C, Figure S3B), differential expression analysis (Figure 2D), and gene ontology enrichment analysis for biologic processes (GOBP, Figure S3C). A similar approach was taken to classify the other cell types comprising the VS TME (Figure S4).

We identified clusters associated with myelination (*e.g.*, *PRX, NCMAP*), hypoxia (*e.g.*, *VEGFA*, *HILDPA*), cell stress (*e.g.*, *JUNB, FOSB*), and interferon-response (*e.g.*, *ISG15, IFIT1*). Two clusters of cells expressed core markers of nmSC identity, including *NRXN1*, *SCN7A*, and *NCAM1*, and largely lacked expression of the other VS-SC clusters (“core”). Interestingly, we noted cells enriched for genes associated with MHC class II antigen presentation (*e.g.*, *CD74, HLA-DRB1*), consistent with SCs in the post-nerve injury setting, which are known to upregulate the antigen-presenting machinery to recruit circulating immune cells and promote their proliferation^33^. Furthermore, two clusters had increased expression of *NGFR, RUNX2*, *SPP1,* and *GAP43*, all of which are upregulated in the setting of peripheral nerve injury (“repair-like”)^34–37^.

Prior studies of VS have suggested that tumorigenic SCs adopt a de-differentiated, immature SC phenotype, while others have suggested that VS-SCs resemble “repair Schwann cells” in the setting of an acute nerve injury^38^. To better understand the phenotypes of VS-SC, we used transcriptional signatures from murine Schwann cells reported in scRNA-seq analyses of peripheral nerves in multiple contexts, including steady-state adult, early development, and post-injury^23,24,26^. Scoring the VS-SCs for each of these signatures indicated that VS-SCs most closely resemble SCs after peripheral nerve injury (Figure 2E). Interestingly, VS-SCs scored low for cycling SC markers seen in these settings. Together, these findings suggest that VS-SC downregulate myelination-associated genes, upregulate gene expression programs that promote nerve repair and immune cell recruitment, and largely remain in a non-proliferative state.

### VS TME immune cells are disproportionately cycling

The observation that VS-SCs do not strongly express markers of proliferation motivated us to return to our analysis of the broader cell type composition of the VS TME, in which we observed a distinct cluster of cells that was driven by cell cycle marker expression (Figure 1C). After assigning these cells to the VS cell type they most closely resembled, we found that VS-SC and stromal cells were underrepresented whereas immune cells were overrepresented in the cycling cell cluster (Chi-squared test, *p* < 0.001; Figure 3A). Next, we turned our attention to all cells across the entire dataset, excluding the cycling cell cluster. We scored each cell type for cell cycle markers and found that immune cells collectively scored higher for both S-Phase and G2M-Phase markers (ANOVA *p* < 0.001; Figure 3B). To validate these observations, we performed immunohistochemical staining of the same tumors used for scRNA-seq. We used CD45 to identify immune cells and Ki67 to identify cycling cells (Figure 3C). Consistent with our scRNA-seq analyses, we found that a higher proportion of CD45 positive cells were Ki67 positive than CD45 negative cells (Figure 3D). Together, these findings suggested that immune cells in the VS TME are disproportionately proliferative and therefore may play a vital role in tumor progression.

**Figure 3.**
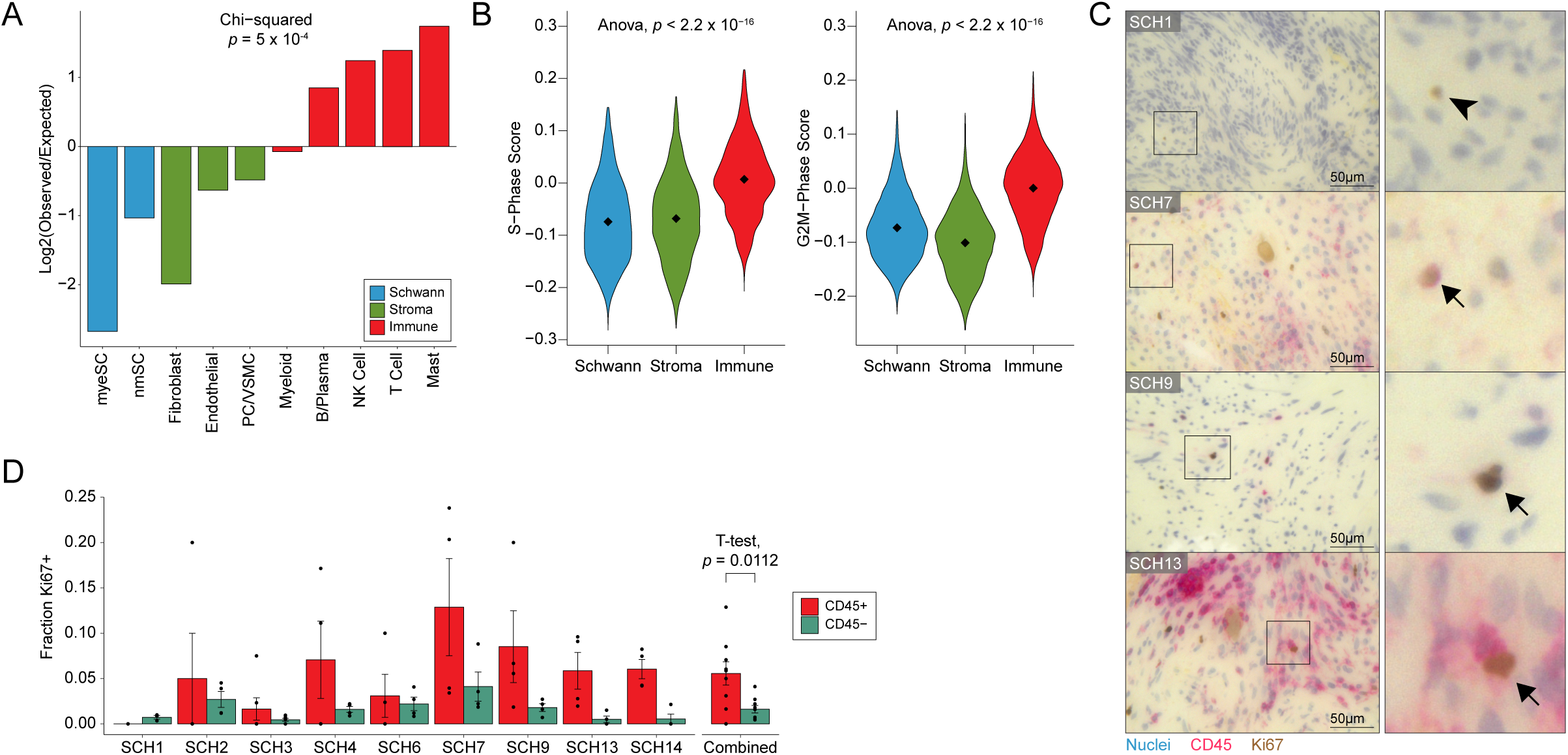
Immune cells are disproportionately cycling in the VS TME. (A) Cycling cells (Figure 1C) were scored based on gene signatures of all other cell types in the VS TME (*e.g.*, nmSC, T cells, *etc.*) and assigned to the cell type for which they scored highest. Frequencies of each cell type observed in this cluster were compared to expected rates. (B) Violin plots of G2M and S-phase scores for Schwann, stromal, and immune cells. (C) Double-stain IHC of representative high-power field (HPF) from VS tumor FFPE samples. Cycling cells are labeled Ki67 and immune cells are labeled with CD45. Arrowhead indicates a representative CD45-Ki67+ cell. Arrows indicate representative CD45+Ki67+ cells. (D) Barplot showing the fraction of CD45+ (red) and CD45- (green) cells that are Ki67+ within available samples (left) and averaged across all samples (right). Error bars on left show standard error for quantification of each group across 4 HPF. Error bars on the right represent standard error of mean measurements across samples.

### VS tumors enriched for nerve injury-related subtypes are associated with increased myeloid cell infiltrate

We next sought to characterize the degree to which VS-SC subtypes varied across samples (*i.e.*, *inter*-tumoral heterogeneity). We assigned subtype scores to each sample by first scoring all VS-SCs for each meta-cluster signature and then taking the mean for each signature. Unsupervised hierarchical clustering of these sample scores revealed two groups of tumors, one enriched for repair-like and MHC II signatures (“Injury-like”) and the other enriched for the core signature (“nmSC Core”) (Figure 4A). These groups differed most by their expression of the repair-like, MHC II, and core programs (Figure 4B; multiple comparisons corrected for with BH method, FDR < 0.2). Interestingly, we found that both the repair-like (*R* = 0.77, *p* < 0.05) and MHC II (*R* = 0.61, *p* < 0.05) scores were associated with an increased fraction of myeloid cells (Figures 4C). The core meta-signature scores did not correlate with degree of myeloid infiltrate. These findings suggest that the VS can be broadly divided into two groups – Injury-like VS and nmSC Core VS – based on the composition of their TME.

**Figure 4.**
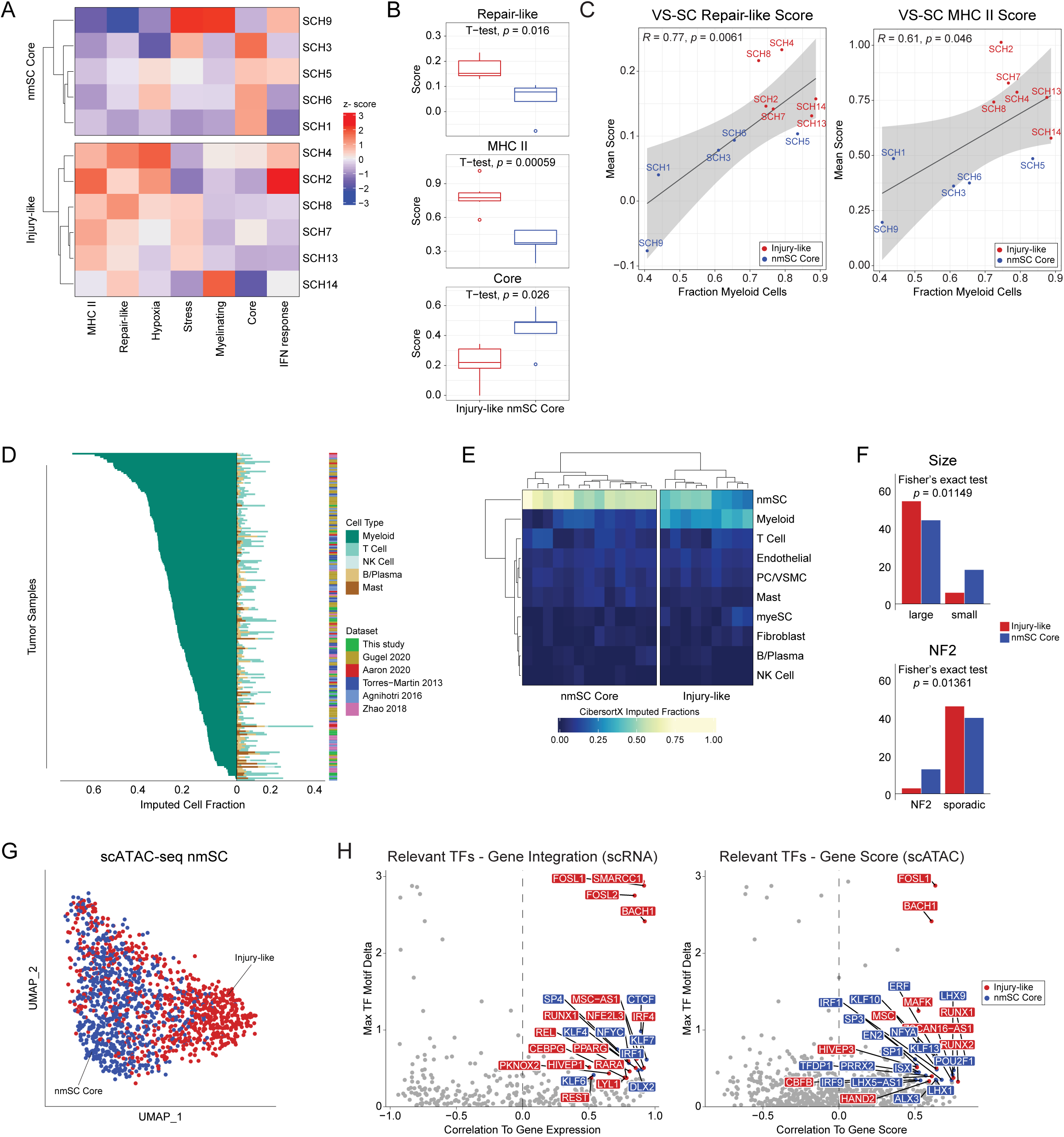
Injury-like VS tumors are associated with increased myeloid cell infiltrate. (A) Heatmap displaying results of hierarchical clustering of VS-SC subtype mean signature scores shows two distinct groups of tumors (“Injury-like” and “nmSC Core”). (B) Box-and-whisker plot comparing mean scores of repair-like, MHC II, and Core signatures in Injury-like and nmSC Core tumors. Two-sided t-testing was performed with correction for multiple comparisons via BH method with FDR of 0.2. (C) Scatterplots demonstrate strong correlation of mean repair-like (left) and MHC II (right) scores with fraction of myeloid cells across samples. (D) Barplot of imputed cell-type fractions from 175 VS tumors shows high variability in degree of myeloid cell composition. Mac, Myeloid; TC, T cell; NK, NK cell; BC, B cell; Mast, Mast cell. (E) Representative heatmap demonstrating classification of our cohort of 22 VS tumors into Injury-like and nmSC Core categories based on hierarchical clustering of imputed cell fractions. Remaining results shown in Figure S5B-F. (F) Barplots showing number of tumor samples classified as Injury-like or nmSC Core and clinically classified by size (n = 122) and NF2-syndrome status (n = 89). (G) UMAP of all VS-SC from the scATAC-seq dataset with cells colored based on the type of VS, Injury-like (red) and nmSC Core (blue), from which they arose as determined by clustering in (A). (H) Scatter plot depicting transcription factor (TF) motif deviation delta between Injury-like and nmSC Core VS-SC and correlation to gene expression (left) and gene score based on accessibility (right). Relevant TFs (correlation > 0.5, adjusted *p* < 0.01 and max delta > 75th percentile of all max deltas) are labeled and colored.

### VS-associated myeloid cells have properties of tumor-associated macrophages and acute inflammatory cells

Since myeloid cells were the most abundant immune cell type in our dataset and therefore might play a role in the pathogenesis of VS, we sought to better characterize the diversity of their functional phenotypes. Given their lack of discrete states, as has been observed in other scRNA-seq studies of human tumors^39^, we utilized a previously described implementation of non-negative matrix factorization (NMF) to identify gene expression programs that recurred across samples (*i.e.,* “meta-programs” (MP))^40^. Using this approach, we identified 69 distinct gene expression programs across patients, of which eight MPs exhibited similar expression across patient samples (Figure S4E-F). Each MP was then annotated according to its functional enrichment. We used gene signatures from recently published pan-cancer and pan-tissue scRNA-seq atlases of myeloid cell phenotypes to evaluate the VS myeloid MP signatures in the context of these integrative resources^39,41^. As expected, we saw marked overlap between the VS myeloid inflammatory MP and pan-cancer M1 signature, the VS angiogenic MP and pan-cancer signatures, and the VS phagocytic MP and pan-cancer signatures (Figure S4G). The pan-cancer M2 signature was less specific, with M2-associated genes expressed across several VS myeloid MPs (*e.g.,* phagocytic, angiogenic, migratory, and granulocytic). This is consistent with more recent observations that macrophages take on a variety of transcriptional states *in vivo* beyond the traditional M1/M2 states^42^. Interestingly, when looking at pan-tissue signatures comparing cancer and inflammatory associated monocytes and macrophages, some VS myeloid cells (*e.g.*, granulocytic, angiogenic, and inflammatory) expressed markers associated with the *inflammatory* monocytic signature while others (*e.g.*, phagocytic, migratory, and oxidative phosphorylation) expressed *cancer* monocyte/macrophage signature genes (Figure S4H). Our analysis suggests that many VS myeloid cells are monocytic in origin with pro-inflammatory, M1-like signatures, while other subsets appear to adopt a spectrum of phenotypes resembling M2-like macrophages.

### Myeloid cell infiltration varies across tumors and is associated with tumor size

To assess the cellular composition of the TME in a larger cohort of patients, we used previously described deconvolution methods on VS tumors characterized with bulk transcriptomic approaches (*i.e.*, RNA-seq and expression microarray)^43^. Using our scRNA-seq gene expression data to define a cell-type signature matrix, we performed digital cytometry using CIBERSORTx on a cohort of 22 newly sequenced tumors combined with bulk transcriptomic data (153 tumors) from previously published reports^27,29–31,44^. Interestingly, we noticed a marked variability in the proportion of immune cells across tumors (Figure 4D). Furthermore, increasing immune cell infiltrate was strongly correlated with the imputed fraction of myeloid cells (*R* = 0.93, *p* < 2.2e^−16^) and only weakly correlated with the fraction of T cells (*R* = 0.26, *p* = 0.00021; Figure S5A), suggesting that variability in immune cell composition is primarily driven by the fraction of myeloid cells. Inversely, the fraction of nmSC was anti-correlated with the fraction of immune cells (*R* = −0.8, *p* < 2.2e^−16^; Figure S5A).

Next, we performed unsupervised hierarchical clustering of the imputed cell fractions from each cohort of bulk expression samples. We found that each dataset could be classified into two distinct cohorts of tumors. One group was characterized by a lower proportion of nmSCs and high myeloid cell infiltrate, reminiscent of the Injury-like VSs in the scRNA-seq analysis, which we labeled “Injury-like”. The other group was characterized by a predominance of nmSCs and low imputed fractions for all other cell types including macrophages, which we labeled “nmSC Core” (Figure 4E, Figure S5B-F). We then assessed whether the Injury-like and nmSC Core cohorts were associated with any clinical parameters of interest. Notably, the nmSC Core tumor group was overrepresented in NF2 syndrome-associated tumors (Figure 4F, Fisher’s exact test, *p* = 0.01149). Furthermore, large tumors (≥ 2 cm in greatest axial dimension or Hannover Scale ≥ 3a) were disproportionately associated with the Injury-like cohort, while small tumors were disproportionately classified as nmSC Core (Figure 4F, Fisher’s exact test, *p* = 0.01361). Comparison of other clinical parameters of interest (prior radiation, hearing loss, tinnitus, vertigo, and tumor consistency) did not reveal any significant associations (data not shown). Thus, across a large cohort of patients, the Injury-like tumor composition is associated with larger tumor size.

### Analysis of chromatin accessibility in Injury-like VS-SC identifies TFs enriched in peripheral nerve injury

Given that Injury-like and nmSC Core VS-SCs differ transcriptionally, we wanted to characterize how these cells might differ epigenetically. We therefore turned our attention to the VS-SCs in the scATAC-seq dataset, which was comprised of three Injury-like and three nmSC Core tumors based on scRNA-seq analysis (Figure 4A). Indeed, after selecting scATAC-seq VS-SCs, and assigning them to either Injury-like or nmSC Core groups based on the tumor from which they were derived, we observed that the Injury-like and nmSC Core cells were distributed differently across UMAP space (Figure 4G). Accordingly, analysis of differentially accessible peaks (DAPs) identified 5616 statistically significant marker peaks with Log2FC ≥ 2 differentiating the two groups of VS-SCs (Figure S6A-B), further suggesting that these two groups of VS-SCs differ from each other significantly at the epigenetic level. Next, we performed TF motif enrichment analysis on a per-cell level based on accessibility of TF binding sites from CIS-BP. We then identified relevant TFs, defined as TFs with gene expression (either inferred from scATAC-seq data or measured from scRNA-seq data) that is positively correlated with increased accessibility of their motif, for Injury-like and nmSC Core SCs (examples of relevant TFs are shown in Figure S6B). Because of the correlation between motif accessibility and associated TF expression, these TFs may be most critical to defining cell state. Indeed, we identified several enriched TF motifs with corresponding increased TF expression among Injury-like (*e.g.*, *BACH1*, *SMARCC1*, *FOSL1*, *FOSL2, RUNX2*) and nmSC Core (*e.g.*, *CTCF*, *NFYC*, *KLF7*) SCs (Figure 4H). Interestingly, many Injury-like TFs have been strongly implicated in the normal SC response to nerve injury^45–48^. For example, an increase in both FOSL2 binding motifs and *FOSL2* gene expression have been found in repair SCs^45^, reminiscent of the repair-like expression profile found in Injury-like VS. In contrast, CTCF was found to be critical for SC differentiation into myelinating SCs, the most mature SC state, consistent with the decreased repair-like expression profile in nmSC Core VSs^47^.

### Injury-like VS-SCs secrete ligands known to promote myeloid cell migration and proliferation

We next sought to characterize the signaling pathways by which VS tumor cells might communicate with other cell populations in the VS TME in Injury-like and nmSC Core tumors. We first focused on tumor-wide patterns of intercellular communication. We inferred network-wide ligand-receptor interactions using CellChat^49^ and found that Injury-like tumors had a higher total number of inferred intercellular interactions and overall higher imputed interaction strength, largely driven by stromal and SC interactions (Figure S6C).

Next, we sought to better understand the specific signaling pathways upregulated and downregulated in Injury-like VSs. Notably, *CCL*, *LIGHT, NECTIN, PERIOSTIN, HGF*, *PTN*, and *CSF* signaling pathways had stronger and more abundant interactions in Injury-like tumors (Figure 5A). A relative increase in outgoing *CCL* signals was observed across all cell types in Injury-like tumors except for mast cells and B cells (Figure 5B), with endothelial cells being the primary receiver of these signals via *ACKR1* expression. *ACKR1* encodes the Duffy antigen receptor, which mediates chemokine transcytosis and enhances leukocyte migration and may therefore promote immune cell recruitment in Injury-like VSs^50^. Interestingly, Injury-like fibroblasts and SCs had increased expression of *HGF* and its receptor, *MET*, respectively. Prior work has established *HGF* as a crucial activator of repair Schwann cells in peripheral nerve injury models, suggesting that this signaling may induce the VS-SC states seen in Injury-likeVSs^51^. Lastly, *CSF* signaling distinctly arose from both myeSC and nmSC in Injury-like tumors, with myeloid cells and cycling cells receiving these signals. Both IL-34 and CSF1 are known chemotactic factors for circulating monocytes secreted by SCs, and previous work has shown that both IL-34 and CSF1 are expressed in VSs, with a weak correlation between tumor growth and CSF1 levels described^52,53^. Our results suggest that this signaling is increased in Injury-like tumors.

**Figure 5.**
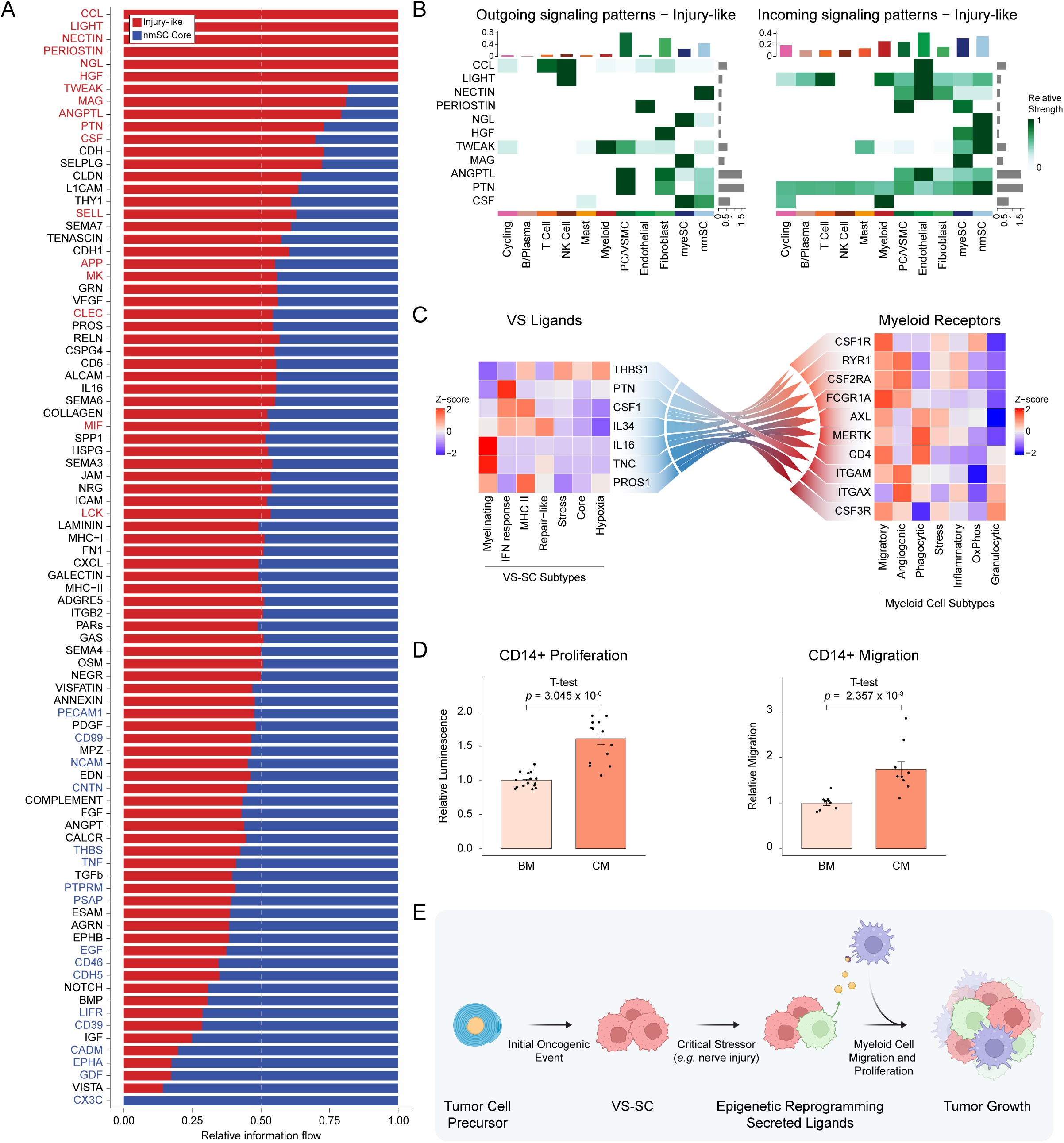
Ligand-receptor interactions in the VS-TME distinguish Injury-like from nmSC Core tumors, and promote myeloid cell proliferation and migration. (A) Bar plot showing the relative information flow of select signaling pathways. Pathway names in red are enriched in Injury-like VS and those in blue are enriched in Core VS. Information flow is defined as the sum of communication probability among all pairs of cell groups in each inferred network. (B) Heatmap displaying the relative interaction strength of signaling pathways enriched in Injury-like VS. Top barplots show summative contribution of individual cell types. Side barplots show summative contribution of a given pathway to the inferred communication network. (C) Heatmap showing relative expression of VS-SC ligands (left) with receptors expressed on myeloid cells (right). (D) Barplots showing relative transwell migration (left) and proliferation at 48 hours (right) of CD14+ monocytes from healthy donors in conditioned media (CM) and base media (BM) from immortalized human Schwann cells (HSC). Each bar represents the normalized mean of all technical replicates (n = 3 per migration assay, n = 5 per proliferation assay) across biological replicates (n = 3). (E) Model of Injury-like VS. VS-SC undergo a critical stressor that triggers subpopulations to adopt repair-like and antigen presenting states. Myeloid cells are recruited to the VS TME and proliferate locally, leading to tumor progression.

Given the abundance of myeloid cells in Injury-like VS, we sought to further characterize VS-SC to myeloid signaling at the cell subtype level. We sought to identify secreted ligands that were 1) strongly expressed by VS-SC in the scRNA-seq data, 2) differentially expressed in tumors relative to healthy nerve controls in the bulk expression data, 3) and had cognate receptors expressed in the VS myeloid cells. Our search identified seven candidate ligands with 10 predicted receptors (Figure 5C). Of note, *IL34* was expressed by repair-like SCs and MHC II SCs, which also highly expressed *CSF1*, with the cognate receptor *CSF1R* most strongly expressed in migratory myeloid cells. We therefore hypothesized that VS-SCs promote myeloid cell migration and proliferation. To test this hypothesis, we applied conditioned media from a previously utilized cell line model of schwannoma (immortalized human Schwann cells; HSC) to human CD14+ peripheral blood monocytes^27^. We found that conditioned media from the schwannoma line promoted the migration and proliferation of monocytes *in vitro*, suggesting that secreted VS-SC factors may influence both processes (Figure 5D). Together these findings suggest that VS-SCs secrete ligands that recruit monocytes and drive their proliferation, potentially contributing to the progression of VS (see model in Figure 5E).

## DISCUSSION

The fundamental factors driving VS tumor progression and unfavorable clinical outcomes remain poorly understood, and consequently, effective medical therapies to limit VS growth remain elusive. Our single-cell multi-omic analysis of sporadic VS represents an important step in understanding the *intra-* and *inter-*tumoral heterogeneity underlying their pathogenesis and progression. Among our key findings is an unexpected diversity within the SC compartment of these tumors. Consistent with prior reports, the majority of VS-SCs are characterized by loss of the myelinating phenotype^38^. Furthermore, using transcriptional signatures derived from the peripheral nerves of mice under steady state, post-injury, and developmental conditions, we found that VS-SCs most resemble SCs in the setting of peripheral nerve injury, with subpopulations of VS-SC adopting transcriptional states similar to repair-type SCs. Interestingly, we noted that, in select tumors, enrichment of repair-like VS-SCs correlated with VS-SCs that express the MHC class II antigen presentation machinery. Furthermore, this group of tumors also had disproportionately higher fractions of cells of myeloid lineage (*e.g.*, monocytes and macrophages) comprising the TME. In the setting of peripheral nerve injury, SCs are believed to be the initial recruiters of monocytes and macrophages, which then contribute to breakdown of myelin and recruitment of additional leukocytes^54^. Accordingly, our findings reveal that the TME of Injury-like VSs resembles the cellular microenvironment of a peripheral nerve in the initial days after injury.

In contrast to damaged peripheral nerves, where SCs proliferate along the trajectory of regenerating axons, we observed low proliferative capacity among VS-SCs in our data, which is consistent with the typical slow growth of these lesions^55^. Interestingly, we found that infiltrating immune cells expressed markers of cell cycle progression at a higher rate than VS-SC or VS stromal cells, which suggests that cues within the VS TME promote this immune cell turnover and renewal. Our findings are consistent with a prior immunohistochemical study of VS tumors with sudden growth, which found that tumor-associated macrophages (TAM) comprised 50-70% of all proliferating cells *in situ*^56^. Thus, our analysis extends on these findings and converges on the overarching principle that myeloid cell proliferation and infiltration may be key cell biological processes that underlie tumor growth.

In our deconvolution analysis of 175 tumors characterized by bulk expression sequencing, we found that Injury-like tumors were associated with larger tumor size. The variable presence of TAMs in the VS TME has been previously described, but their role in VS pathogenesis and their functional phenotypes have been poorly characterized^52,56,57^. For example, increased presence of macrophage markers on histology has been associated with tumor growth, poor post-operative facial nerve outcomes, and poor pre-operative hearing^56,58,59^. Other reports have suggested that an inflammatory dimension of VSs may contribute to adverse outcomes in these patients and have served as the basis for ongoing trials evaluating the potential of aspirin to mitigate sudden tumor growth^60^. Interestingly, among this broad cohort of patients, NF2-associated VS tumors were almost exclusively low in macrophage infiltrate. Why these lesions harbor fewer infiltrating immune cells remains an important question, as our cohort of patient samples characterized by scRNA-seq did not include any syndromic NF2 patient tumors. Future work characterizing both sporadic and syndromic VS will help elucidate the differences in microenvironmental cues that promote myeloid cell recruitment in specific tumors.

Given that Injury-like VSs may be associated with worse patient outcomes, we sought to characterize the transcriptional regulation and cell-to-cell signaling of these tumors relative to nmSC Core VSs to identify potentially novel therapeutic targets. We found that nmSCs from Injury-like and nmSC Core tumors bear different epigenetic profiles. Furthermore, we identified several relevant TFs that not only have accessible motifs in both Injury-like and nmSC Core cells but also demonstrated increased gene expression of the relevant TF in the respective VS-SC groups (*e.g., RUNX1*, *FOSL1*, *FOSL2*, *etc.*). Regarding cell-to-cell signaling, while there were pathways more highly expressed in Injury-like tumors (*e.g., CCL*, *MIF*, *etc.*), CSF signaling appeared to be specific between VS-SC and myeloid cells. This signaling axis is seen in inflammatory neuropathies, and our results suggest its role may extend to VS tumor progression ^53,61^. Experiments using an *in vitro* VS model and healthy donor CD14+ monocytes further support the hypothesis that VS-SCs promote monocyte migration and proliferation. Taken together, our findings uncover potential pathophysiological mechanisms that may drive tumor growth and require major investigation, including future pre-clinical work to screen regulatory transcription factors and/or receptor-ligand pathways for their effects on tumor behavior.

There are several limitations of this study. Patients in our scRNA-seq cohort were limited to sporadic VS, and our findings pertaining to the TME composition and SC states may not be generalizable to patients with schwannoma of other sites or patients with syndromic NF2- associated tumors. In addition, our patient cohort was restricted to patients who underwent surgery, and thus we were unable to characterize small, asymptomatic tumors since such lesions are routinely observed radiographically or treated with stereotactic radiosurgery.

In summary, our work provides important insights into VS biology as well as a detailed transcriptomic and epigenetic single cell atlas of the Schwann, stromal, and immune cells that comprise the VS TME. Our analysis suggests that VSs can be categorized based on nerve Injury-like VS-SC gene expression programs and associated myeloid cell infiltrate. Furthermore, Injury-like tumors appear to be associated with larger tumor size, and chemokines secreted by VS-SCs may recruit circulating monocytes. These findings uncover previously undescribed mechanisms of pathogenesis and tumor progression in VS and suggest novel biomarkers and therapeutic targets to be explored in future studies.

## MATERIALS AND METHODS

### Human tumor specimens

Patient samples used for scRNA-seq and scATAC-seq were all derived from patients treated at Barnes-Jewish Hospital (St. Louis, MO, USA). All patients provided written informed consent to participate in the study following Institutional Review Board Approval (Protocol #201111001, #201103136, and #201409046). Patient characteristics are summarized in Figure 1B and Table S1. Tumor samples used for bulk RNA-seq analysis consisted of paraffin-embedded tissue from 22 VS patients treated at Baylor College of Medicine (BCM; Houston, TX, USA) (Table S2). All patients provided written informed consent, and tumor tissues were collected under an institutional review board (IRB)-approved protocol at BCM by the Human Tissue Acquisition and Pathology Core (Protocol H-14435). All schwannomas were reviewed by a board-certified neuropathologist according the 2016 WHO guidelines. Raw data from previously published studies were obtained as follows: RNA-seq and expression microarray data that were publicly available were downloaded (GSE39645^30^, GSE141801^29^, GSE108524^31^, EGA00001001886^27^); data from *Aaron et al*^44^ were kindly shared upon request. Clinical annotations accompanying the sample data from *Torres-Marin et al*^30^ were also kindly shared upon request.

### Fresh tumor dissociation

Samples processed for scRNA-seq and scATAC-seq were collected at the time of surgical resection and immediately processed. Tumor samples were minced and dissociated using the Human Tumor Dissociation Kit (Miltenyi Biotech, Bergisch Gladbach, Germany) per manufacturer guidelines. The dissociated cell suspensions were then passed through 40µm filter, pelleted through centrifugation, and resuspended in AutoMACS Rinsing Solution with 0.5% bovine serum albumin (BSA; Miltenyi Biotech). Red blood cell lysis was performed on all samples with Gibco ACK Lysing Buffer (ThermoFisher Scientific, Waltham Massachusetts, US) and was followed by debris removal via density gradient when necessary (Debris Removal Solution, Miltenyi Biotech, Bergisch Gladbach, Germany). Cell viability was confirmed to be > 80% using 0.4% Trypan Blue staining (Invitrogen, catalog #T10282) and manual counting with a hemocytometer. For samples in which scATAC-seq was additionally performed, nuclei isolation was performed according to the 10X Demonstrated Protocol “Nuclei Isolation for Single Cell ATAC Sequencing” (Rev D).

### Tumor nuclei isolation for scRNA-seq

Fresh frozen samples used for scRNA-seq were collected at the time of surgical resection and frozen in OCT compound embedding media (Tissue-Tek, Torrance, California) on a pre-chilled aluminum block resting on dry ice, and stored at −80 °C. Tissue scrolls were cut at 30 µm using a Cryostat (50-100 scrolls were cut per sample, depending on the tissue size) and maintained at −80°C until the time of nuclei isolation. Lysis buffer (consisting of Tris-HCl, NaCl, MgCl_2_, Nonidet P40 Substitute, 0.1M DTT, RNase inhibitor, and nuclease free water) was added to the tissue scrolls, which were homogenized using a Pellet Pestle while on ice. Additional lysis buffer was then added, and the mixture was incubated on ice for 5 minutes. The suspension was passed through a 70 µm strainer and centrifuged before being washed with a solution of PBS with 1% BSA and 1U/µl Rnase inhibitor, incubated on ice for 5 minutes, centrifuged, and resuspended in 1ml PBS with 1% BSA and 1 U/µl Rnase inhibitor. The nuclei were then labeled with DRAQ5 (Thermo Scientific, catalog #62251) and selected using FACS sorting performed by the Siteman Flow Cytometry Core before being carried forward for single nuclei library creation.

### scRNA-seq library preparation and sequencing

Single cell and single nuclei suspensions were processed using 10X Chromium Next GEM Single Cell 3’ Reagent Kits v3.1 (10X Genomics, Pleasanton, CA) per manufacturer protocols. Briefly, cells were added onto the 10X Next GEM Chip G to form Gel Bead-in-Emulsions (GEMs) in the Chromium instrument followed by cell lysis, barcoding, cDNA amplification, fragmentation, adaptor ligation, and sample indexed library amplification. Completed gene expression libraries were sequenced on Illumina NovaSeq S4 flow cells at a target depth of 50,000 read pairs per cell. Single cell RNA and single nucleus RNA sequencing reads were aligned to human reference GRCh38 v2020-A from 10x Genomics using the 10x Genomics Cellranger-4.0.0 and Cellranger-6.0.0 (include-introns flag set to true) pipelines, respectively. Sequencing quality control metrics are listed in Table S3.

### snATAC-seq library preparation and sequencing

snATAC-seq libraries were prepared using the 10X Chromium Next GEM Single Cell ATAC Reagent Kits v1.1 (10X Genomics) according to the manufacturer’s protocols. In brief, nuclei were incubated in a transposition mixture including a transposase to fragment open chromatin regions. Transposed nuclei were then loaded onto the 10X Next GEM Chip H to generate GEMs, followed by sample indexed library amplification. snATAC-seq libraries were sequenced in Illumina NovaSeq S1 flow cells at a target depth of 250M total read pairs per sample. The resulting FASTQ files were aligned to GRCh38 v2020-A using the 10x Genomics Cellranger ATAC-1.2.0 count function.

### scRNA-seq/snRNA-seq data preprocessing

Ambient RNA removal and empty droplet calling was performed using CellBender^62^. Samples were processed individually and iteratively with adjustment of the parameters to achieve optimal learning curves and barcode rank plots for each sample. Final parameters used are listed in Table S4. CellBender outputs consisting of counts matrices adjusted for ambient RNA and excluding empty droplets were then preprocessed for doublet calling using Scrublet^63^ and ScanPy^64^ as follows: a) Cells with < 500 genes were excluded; b) Genes not expressed in at least 0.1% of cells were excluded; c) Percent mitochondrial counts was computed for each cell, Leiden clustering performed, and cells with percent mitochondrial counts greater than 2 standard deviations from their respective cluster mean percent mitochondrial counts were removed. Samples were then processed individually and iteratively, varying the n-neighbors and expected_doublet_rate and choosing the values for each that resulted in a bimodal simulated doublet histogram with a bimodal curve fit *R* > 0.85 and the fraction of the second Gaussian less than or equal to the 99th percentile of the first.

The filtered gene expression matrix was then processed and analyzed by using Seurat v4.0.0^65^. To filter low-quality cells, we first removed cells for which less than 1000 genes were detected or cells that contained greater than 20% of genes from the mitochondrial genome. We included genes with ≥5 UMI in at least 10 cells for downstream analysis.

### scATAC-seq data preprocessing and clustering analysis

scATAC-seq preprocessing and analysis was performed using ArchR 1.0.1 as detailed in the ArchR manual^66^. Briefly, nuclei with a TSS < 10 and with < 1000 fragments were excluded. Doublets were identified and removed using the ArchR addDoubletScores and filterDoublets functions with filterRatio = 1.5, DoubletScore ≤ 50. Dimensional reduction was performed using the addIterativeLSI function and default ArchR values of sampleCells = 10000, n.start = 10 and varFeatures = 15000. Next, the addClusters function was used for cell clustering and the addGeneIntegrationMatrix function was used to perform unconstrained cross-platform linkage of scATAC-seq cells with snATAC-seq cells from the scRNA-seq atlas without single nucleus samples. scATAC-seq clusters were then labeled with a cell identity by creating a confusion matrix between scATAC-seq clusters and cell identities from linked scRNA-seq cells and assigning each cluster the identity of the greatest proportion of linked scRNA-seq cells in that cluster (Figure S2E).

### Multiple sample integration with reciprocal principal component analysis

To overcome batch effects related to freshly dissociated samples and nuclei isolated from fresh frozen samples, Seurat’s reciprocal principal component analysis (RPCA) was used to integrate the scRNA-seq datasets^67^. In brief, a SeuratObject was generated for each sample. Each sample was then normalized using Seurat’s ‘NormalizeData’ function. ‘FindVariableFeatures’ was used to identify 3000 variable features in each sample. Integration features were selected using ‘SelectIntegrationFeatures’ (nfeatures = 3000). ‘FindIntegrationAnchors’ was used to perform RPCA integration (by sample) in Seurat. The data was integrated using ‘IntegrateData’ with k-nearest neighbors (k.weight) set to 50; integrated values were returned for all genes in the SeuratObject. The integrated RPCA object was further scaled using ‘ScaleData’ function and was projected on the UMAP with 30 principal components. Graph-based clustering was performed (resolution = 0.5) on the integrated object. Differentially expressed genes were calculated for the clusters of “integrated Assay” on the “RNA Assay” using the ‘FindAllMarkers’ function with only.pos = T (*i.e.*, only for upregulated genes). Only significant (p.adj ≤ 0.05) DEGs were used in further analysis.

### Gene signature scoring and cell type assignments

To corroborate our cell type labels, we used the top 30 differentially expressed genes (DEGs) from each peripheral nerve cell-type cluster as defined by the original authors from each study to score each cell in our VS dataset. The mean score of each signature was calculated for each VS TME cluster using the Seurat AddModuleScore function (Figure S1B). To assess the consistency of peripheral nerve cell-type scores across studies, we assigned meta-signatures for similarly labeled cell clusters within and across the mouse nerve studies (*e.g.*, “Schwann cells” from Carr et al and “Nm-SCs” from Yim et al were assigned the meta-label “Schwann”) and computed the mean score of all cluster scores per meta-signatures (Figure 1F).

### Inferred copy number alteration analysis

CONICSmat (0.0.0.1) was used for single cell CNV analysis^28^. Putative normal and tumor cells were selected based on initial cell type assignment. All the Schwann cells (nmSC and myeSC) were assumed to be tumor cells and the rest of the cells in the immune and stromal component were assumed to be normal cells for input in CONICSmat. Relative count normalization was performed in Seurat (4.0.0) with scale factor of 10^5^ and log2(CPM/10+1) transformation was performed on the resulting matrix. A normalization factor was calculated for each column in the expression matrix using ‘calcNormFactors’ function in CONICSmat. A two-component Gaussian Mixture Model was estimated for the log2(CPM/10+1) expression matrix. Fit-data plots and z-score heatmaps were generated using ‘plotAll’ function in CONICSmat. The fit-data plots were assessed manually for each sample and CNV alteration for its validity. Only the CNVs which showed clear amplification/deletion were chosen for each sample based on the fit-data plots, BIC score (≥ 50) and z-score heatmaps. For each significant CNV alteration, specific clusters that showed the alteration were selected from the z-score heatmaps. For each CNV alteration, a Fisher exact test was used to confirm enrichment of that CNV alteration in the Schwann cells in comparison to other cells.

### Comparison of nmSC and myeSC gene signatures of VS tumor samples to normal nerve

Microarray datasets (GSE141801, GSE108524 and GSE39645) were downloaded using GEOquery’s (v2.58.0) ‘getGEO’ function. Biobase’s (v2.50.0) ‘exprs’ function was used to extract the microarray eSets (expression data from sets) object and log2 normalization was performed. The design matrix for a particular microarray dataset was constructed to compare the type of tissue (*i.e.*, ‘Normal-nerve’ *vs.* ‘schwannoma’) using the ‘model.matrix’ function from stats package (v4.0.3). The eSet object was weighted based on the design matrix and a linear model was fit to the data using limma’s (v3.46.0) ‘arrayWeights’ and ‘lmFit’ functions respectively. ‘makeContrasts’ function from limma was used to extract contrasts between ‘control/normal-nerve’ and ‘tumor/schwannoma’ samples. Empirical Bayes statistics were used for differential expression analysis between normal and tumor samples using limma’s ebayes function. The resulting moderated t-statistics were classified into ‘up’, ‘down’ or ‘no change’ using limma’s ‘decideTests’ function. The scaled eSet matrix was further visualized for top 50 differentially expressed single cell markers from both ‘nmSC’ and ‘myeSC’ cells. ComplexHeatmap (v2.11.1) was used to annotate differential expression and normal-tumor groupings.

### VS-SC, stromal, and NK/T cell analysis

Clusters were extracted from the full scRNA-seq dataset and were renormalized and reclustered using Seurat. The subclusters were corrected/integrated using RPCA, as described above. Samples with fewer than 40 cells for a given cell type were excluded. Clusters that were presumed residual doublets (*e.g.,* cells expressing *PTPRC* in the Schwann cell subcluster) or low quality cells (*i.e.*, high ribosomal RNA content) were manually removed and the remaining data were reprocessed, as above. Due to batch effects that were apparent at the subcluster level between the freshly dissociated cells and isolated nuclei from frozen tissue, we performed the primary subtype analysis on the freshly dissociated samples, with the fresh frozen samples serving as a validation dataset (Figure S3D). Gene Ontology Biologic Process Enrichment analysis was performed using the ‘compareCluster’ function from ClusterProfiler (v3.18.1), with the top 25 DEGs of each celltype subclassification, ranked by average Log2FC. VS-SC were scored using the mouse peripheral nerve Schwann cell-specific DEGs as defined by the original study authors’ labels with Seurat’s ‘AddModuleScore’ function.

### Cycling cell analysis

Cells from the scRNA-seq data that clustered by expression of cell cycle markers (“Cycling Cells”, Figure 1C) were subset from the overall dataset and scored by top 30 DEGs of all other broad cell types comprising the VS TME with Seurat’s AddModuleScore function. Cell-type frequencies were scaled to reflect cell numbers of the overall dataset. Chi-square testing was used to compare scaled expected cell-type frequencies with observed cell type frequencies across the entire dataset. Cell cycle phase assignments were made using Seurat’s CellCycleScoring function with Seurat’s included S-phase and G2M phase markers.

FFPE VS specimens from included patients in scRNA-seq analysis were obtained and used to generate a tissue microarray (TMA). The TMA was designed to include four separate 2mm cores from each FFPE block used for pathologic diagnosis at the time of surgery. Tissue arrays were cut into sections (5μm) on positively charged slides. For immunohistochemistry, sections were stained using a Bond RXm autostainer (Leica). Briefly, slides were baked at 65°C for 4hrs and automated software performed dewaxing, rehydration, antigen retrieval, blocking, primary antibody incubation, post primary antibody incubation, detection (DAB) and (RED), and counterstaining using Bond reagents (Leica). Samples were then removed from the machine, dehydrated through ethanols and xylenes, mounted and cover-slipped. Antibodies for Ki67 (Abcam ab16667) and CD45 (Agilent M0701) were diluted 1:200 in Antibody diluent (Leica). Brightfield images of 3-4 high-power field regions (40x) per patient were obtained using a Nikon ECLIPSE Ti2 inverted microscope. Quantification of cell type marker scoring was performed in a semi-quantitative fashion using QuPath-0.3.1. The ‘Positive Cell Detection’ function was used to identify Ki67+ and Ki67-cells using the following parameters: Nucleus Parameters (Requested pixel size 0.5 µm, Background radius 8 µm, Median filter radius 0 µm, Sigma 1.5 µm, Minimum area 10 µm^2^, Maximum area 40 µm^2^), Intensity Parameters (Threshold 0.001, Max background intensity 2), Cell parameters (Cell expansion 0 µm), Intensity threshold parameters (Score compartment “Nucleus: DAB OD Mean”, Single Threshold 1.4976). CD45+ cells were manually annotated. Statistical analysis was performed using a two-sided student’s t-test to compare the means of individual sample means with a significance threshold of *p* < 0.05.

### Classification of scRNA-seq VS-SC as Injury-like and Core

VS-SC obtained via scRNA-seq were subset and, using the top 50 DEGs of each VS-SC subtype based on average log2FC, scored for each of the identified VS-SC subtypes with Seurat’s ‘AddModuleScore’ function. Individual cell scores were averaged across all cells of a given VS-SC subtype across all samples. Sample scores were scaled and samples were hierarchically clustered based on their scaled scores in an unsupervised manner based on Euclidean distance. The highest branchpoint of the dendrogram was used to divide the cohort into two groups, which we ultimately labeled Injury-like and nmSC Core. Mean scores for each VS-SC subtype were compared between Injury-like and Core using a student’s t-test with correction for multiple hypothesis testing using the BH method with an FDR or 20%.

### Myeloid cell analysis

To identify cell states in Myeloid subcluster, non-negative matrix factorization was applied to each sample to identify meta-programs, as previously described^40^. The data was first normalized using CPM normalization and was transformed with log2(CPM+1) transformation. The CPM expression was then centered across each gene by subtracting the average expression of each gene across all cells. All negative values were then transformed to zero. The NMF was computed on the relative expression values with number of factors (K) ranging from 2-9. For each value of K, the top 100 genes (with respect to NMF score) were used to define an expression program. For each sample, we selected “robust” expression programs, which were defined as having an overlap of at least 70% (intra_min = 70) with a program obtained from the same sample using a different value of K. We removed “redundant” programs, which were defined as overlapping another program from the same sample by more than 10% (intra_max = 10). The programs were filtered based on their similarity to programs of other samples (inter_filter = True). Only those programs which had an overlap of at least 20% between programs of two samples were considered (inter_min = 20). To identify MPs across samples, we compared expression programs by hierarchical clustering, using 100 minus the number of overlapping genes as a distance metric. Eight clusters (*i.e.*, MPs) were defined by manual inspection of the hierarchical clustering results. Final MP signatures only included those genes that occurred in 50% of the constitutive programs per cluster. Individual myeloid cells were scored according to these MP signatures using Seurat’s AddModuleScore function, and the cells were assigned to the metaprogram for which they scored most highly. The functional annotation of these metaprograms was done using (1) GO term enrichment (data not shown) and (2) overlap of these metaprogram genes in existing myeloid subtype markers.

### Bulk RNA sequencing, alignment, and preprocessing

Bulk RNA-sequencing of VS was performed by Tempus, Inc. (Chicago, IL, USA), which entailed sending tumor samples along with saliva for processing according to their protocol^68^. RNA-seq reads were then aligned to the GRCh38 assembly with STAR version 2.7.2b (Parameters:--genomeDir Ensembl_GRCh38.fa --genomeLoad NoSharedMemory --outSAMmapqUnique 60 --outSAMunmapped Within KeepPairs --outFilterIntronMotifs RemoveNoncanonicalUnannotated --outSAMstrandField intronMotif --runThreadN 8 --outStd BAM_Unsorted --outSAMtype BAM Unsorted --alignTranscriptsPerReadNmax 100000 --outFilterMismatchNoverLmax 0.1 --sjdbGTFfile Ensembl_GRCh38_genes.gtf > genome_accepted_hits.bam). Gene counts were derived from the number of uniquely aligned unambiguous reads by Picard version 2.6.0. Sequencing performance was assessed for the total number of aligned reads, total number of uniquely aligned reads, and features detected. All gene counts were then imported into the R (3.2.3). Bioconductor (3.2) package EdgeR and TMM normalization size factors were calculated to adjust for samples for differences in library size. The previously published RNA-seq datasets were aligned and processed in an identical manner.

### Deconvolution analysis of bulk expression data

CIBERSORTx was used to build a custom signature reference from the scRNA-seq dataset and impute cell fractions from each of the RNA-seq and microarray expression datasets on a one-by-one basis to avoid confounding batch effects^43^. Default CIBERSORTx parameters for generation of a scRNA-seq reference matrix were used, except for fraction of cells expressing a given gene, which was set to 0 to avoid overly aggressive filtration of genes for generation of the signature matrix given the sparse nature of 10X Chromium derived data. S-mode was used for batch correction during imputation of cell fractions from mixture (*e.g.*, bulk sequencing) data. Unsupervised hierarchical clustering based on Euclidean distance was performed across all samples for each individual bulk expression dataset, and cohorts were grouped into “Injury-like” and “nmSC Core” Cohorts based on the first dendrogram branchpoint. Samples with available clinical data were split by Injury-like/nmSC Core groups and outcomes of interest were compared across these two groups using a Fisher’s exact test.

### scATAC-seq VS-SC analysis

All VS-SC from the scATAC-seq dataset were subset and assigned an identity of Injury-like or nmSC Core based on the classification of the tumor from which they arose by scRNA-seq analysis. Myelinating SC arose predominantly (> 90%) from a single nmSC Core sample and were therefore excluded from further analysis. To reduce biasing by outlier cells when comparing the two groups, cells in the top and bottom 5^th^ percentile for number of fragments, TSS enrichment, and reads in TSS were excluded from further analysis. Approximately 750 cells remained in each of the Injury-like and nmSC Core groups after filtration and were analyzed further. Pseudo-bulk replicates were created using the ArchR addGroupCoverages function with minReplicates = 3, minCells = 100, maxCells = 500, and sampleRatio = 0, and peak calling was performed using MACS2 (2.2.7.1) (https://pypi.org/project/MACS2/) as detailed in the ArchR manual. Per-cell transcription factor motif deviations were added using the addDeviationsMatrix function and motifs annotated using the CIS-BP annotations built in to ArchR. Positive transcription factor regulators were identified using the correlateMatrices function and pairing either the gene score matrix (containing chromosomal accessibility data) or the gene integration matrix (containing gene expression data from linked scRNA-seq cells) with the transcription factor deviations matrix (see ArchR manual for details). Relevant TFs were defined based on default ArchR parameters (correlation > 0.5, adjusted *p* < 0.01 and max delta > 75^th^ percentile of all max deltas).

### Ligand-receptor analysis

Cell-cell communication networks were inferred using the standard CellChat inference and analysis of cell-cell communication workflow CellChat (1.5.0)^49^. In brief, the scRNA-seq was divided into two cohorts (Injury-like and Core), each individual dataset then underwent library size normalization followed by log transformation using *Seurat’s* ‘NormalizeData’ function. The CellChatDB curated database of ligand-receptor interactions was used, over-expressed ligand/receptor genes were identified within each broad cell group (*e.g.*, nmSC, fibroblasts, *etc.*) using the ‘identifyOverExpressedGenes’ function, and then each ligand-receptor interaction were identified using the ‘identifyOverExpressedInteractions’ function. Communication probabilities were calculated for both ligand-receptor pairs and pathway level interactions using the ‘computeCommunProb’ and ‘computeCommunProbPathway’ functions, respectively. The cell-cell communication networks were then summarized using the ‘aggregateNet’ function to determine the number of unique links and overall communication probability. The two communication networks (*i.e.*, Injury-like VS and nmSC Core VS) were compared following the CellChat manual for comparison analysis of multiple datasets. Functions were performed with default parameters unless otherwise stated. Total interactions and interaction strength were determined using the ‘compareInteractions’ function and visualized on a cell-type level as a heatmap using the newVisual_heatmap’ function. Joint manifold learning and classification of the inferred communication networks based on their functional similarity was performed using the ‘computeNetSimilarityPairwise’, ‘netEmbedding’, and ‘netClustering’ functions. Conserved and context-specific signaling pathways for each communication network were compared using the ‘rankNet’ function and a Wilcoxon rank-sum testing was performed with *p* cutoff of 0.05. Cell-type population level signaling was visualized in a heatmap using the ‘netAnalysis_signalingRole_heatmap’ function for those pathways that were most specific to Injury-like tumors (Figure 5A).

Specific interactions between VS-SC and myeloid cells were determined in the following manner. First, we used an extensive, previously described ligand-receptor database to identify potential signaling pairs (*NicheNet*)^69^. We identified ligands expressed in the VS-SC populations with an average Log2FC of 0.5 and expression in at least 5% of VS-SC and with similarly expressed cognate receptors in the myeloid cells. This list was further refined by only including ligand and associated receptor genes that were differentially expressed by tumors relative to normal nerve controls in the expression microarray datasets, as described above. Lastly, the resulting list was filtered to only include those ligands that were known to be secreted molecules by review of the existing literature.

### Cell lines

HSC cells were generously provided by Dr. Gelareh Zadeh and colleagues. They were cultured in DMEM (ThermoFisher Scientific) supplemented with 10% fetal bovine serum (FBS) (Peak Serum, Fort Collins, CO) 1X penicillin-streptomycin (PSG) (ThermoFisher Scientific), and supplemented with 2 µL forskolin (Sigma-Aldrich).

### CD14+ monocyte isolation

Peripheral blood mononuclear cells (PBMC) were obtained from leukocyte reduction system cones that are classified as non-human research under the Washington University Human Research Protection Office. PBMC were isolated using SepMate tubes (StemCell Technologies) and Ficoll-Paque density gradient medium (Fisher Scientific). CD14+ cells were positively selected using anti-CD14-conjugated magnetic microbeads (Miltenyi Biotec).

### Migration assay with conditioned media

Conditioned media (CM) was obtained as follows: HSC cells were plated at a density of 500,000 cells/10cm tissue culture plate in their growth media containing 2.5% FBS. CM was collected at 72 hours after plating, passed through 0.45 µM PES syringe filter (MidSci), and used fresh. Base media (BM) served as a negative control and consisted of growth media for each respective line with 2.5% FBS that was placed in an empty tissue culture plate in parallel to the CM plates, collected and filtered at 72 hours, identically as the CM. 500 µL of CM or BM were placed into wells of a 24-well tissue culture plate. Cell culture inserts (8 µm; Corning) were placed into each well and CD14+ cells were plated above the inserts at a density of 1×10^6^ cells in 250 µL serum-free RPMI 1640 media. Plates were incubated at 37° C for 24 hours. Quantification was performed by manual cell counting of the media in the bottom wells using a hemocytometer. Each condition was performed in triplicate, and the experiment was repeated three times to ensure biologic validity.

### Cell proliferation with conditioned media

CellTitre-Glo (CTG) proliferation assays (Promega) were completed according to manufacturer protocols. Briefly, 1000 CD14+ monocytes were seeded per well in a 96 well plate in 100μL of HSC CM or HSC BM in technical replicates of 5. Cells were lysed on day 0 (one hour after seeding of cells) and day 2 by addition of the CTG reagent followed by measurement of luminescence using the Biotek Cytation 5 (BioTek, Winooski, VT). Luminescence values were adjusted based on 2µM Adenosine triphosphate (ATP) luminescence measured on the same plate for each day and background luminescence was removed. CM and BM were prepared, as above, except that media contained 10% FBS. The experiment was repeated three times to ensure biologic validity.

## ACKNOWLEDGEMENTS

We would like to acknowledge: Gelareh Zadeh and her laboratory for providing HSC cell lines, cell culture methods and sequencing data, Zarko Manojlovic for providing bulk RNA sequencing data, Miguel Torres-Martin for providing clinical data, Travis Law for assistance in implementation of scRNA-seq preprocessing methods, and Raleigh Kladney for immunohistochemistry assistance. Portions of figure 1A and 5E were created with BioRender.com.

## AUTHOR CONTRIBUTIONS

T.F.B. and B.P. contributed equally to the work and A.H.K., S.V.P. and A.A.P. jointly supervised the work. T.F.B. and B.P. performed experiments, data analysis, and manuscript and figure preparation. S.M.K., A.K.Y.Y., and S.P. assisted with data analysis. T.M. and R.D.Z.M. assisted with methods development. All authors contributed to manuscript review/editing.

---

**Figure S1.**
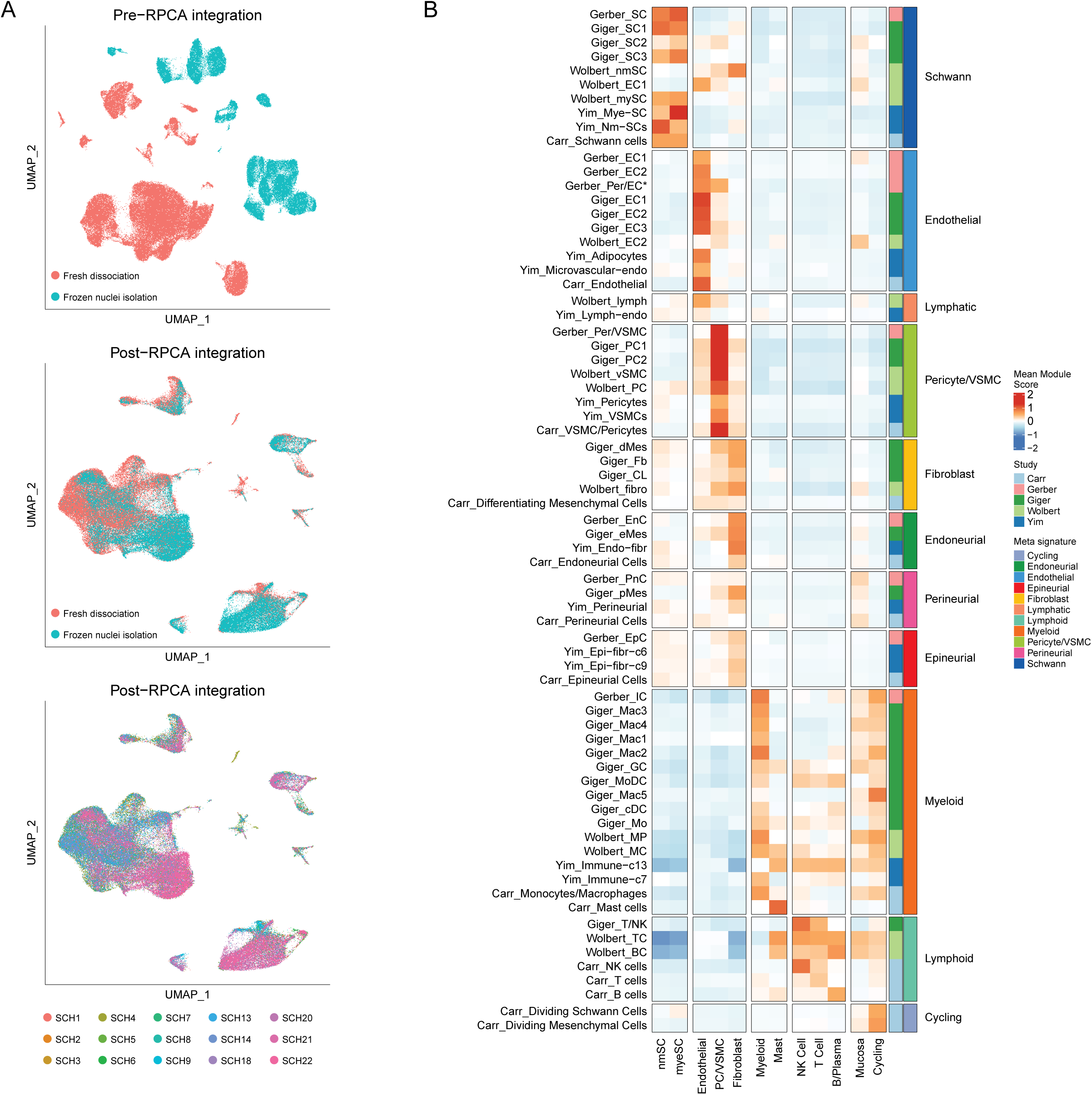
scRNA-seq data integration and VS TME classification by peripheral nerve signatures. (A) UMAP plot of VS scRNA-seq dataset of embeddings pre-RPCA integration (top), post-integration (middle), and colored by individual sample. Batch effects associated with fresh tissue dissociation and nuclei isolation are overcome. (B) Heatmap showing average mouse peripheral nerve cell-type signature score (rows) for each VS cell-type cluster (columns).

**Figure S2.**
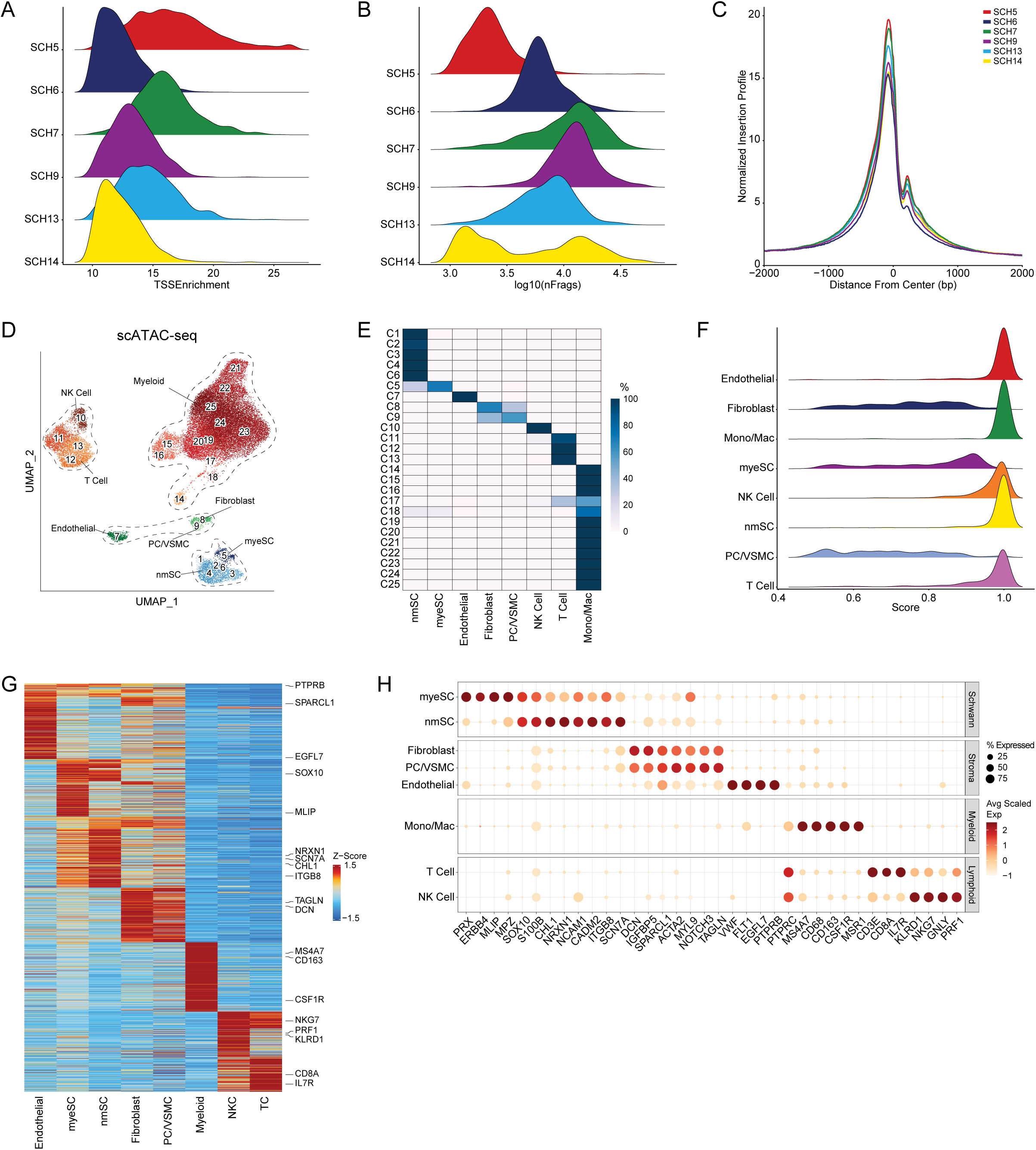
scATAC-seq quality control and cell type assignment. (A) Ridgeplot showing distribution of TSS Enrichment for each tumor after preprocessing (removal of cells TSS Enrichment < 10, number of fragments < 1000, and doublet removal as detailed in text). (B) Ridgeplot showing distribution of number of fragments per cell for each tumor after preprocessing. (C) TSS enrichment profile for each sample shows clear peak in center with smaller shoulder peak to the right, consistent with a well-positioned +1 nucleosome and good quality ATAC-seq data. (D) “Over-clustering” of scATAC-seq data at high resolution identifies 25 cell clusters which were initially labeled using inferred expression from gene accessibility of marker genes of various cell types. NK, natural killer cells; VSMC, vascular smooth muscle cells; nmSC, non-myelinating Schwann cells; myeSC, myelinating Schwann cells. (E) Confusion matrix generated after unconstrained linkage of cells from the scRNA-seq dataset with cells in the scATAC-seq dataset. scATAC-seq cells in most clusters where overwhelmingly linked with just one type of scRNA-seq cell. scATAC-seq clusters were ultimately labeled with the identity of the scRNA-seq cell type that most cells in the cluster were linked to. (F) Ridgeplot displaying distribution of score assigned to each scATAC-seq to scRNA-seq cell linkage by ArchR. (G) Heatmap of marker genes for each cell cluster identified from inferred gene expression based on chromatin accessibility. (H) Dot plot of expression levels of characteristic genes described in the literature for each VS cell subpopulation derived from expression data of scRNA-seq cells linked to scATAC-seq cells in each cluster.

**Figure S3.**
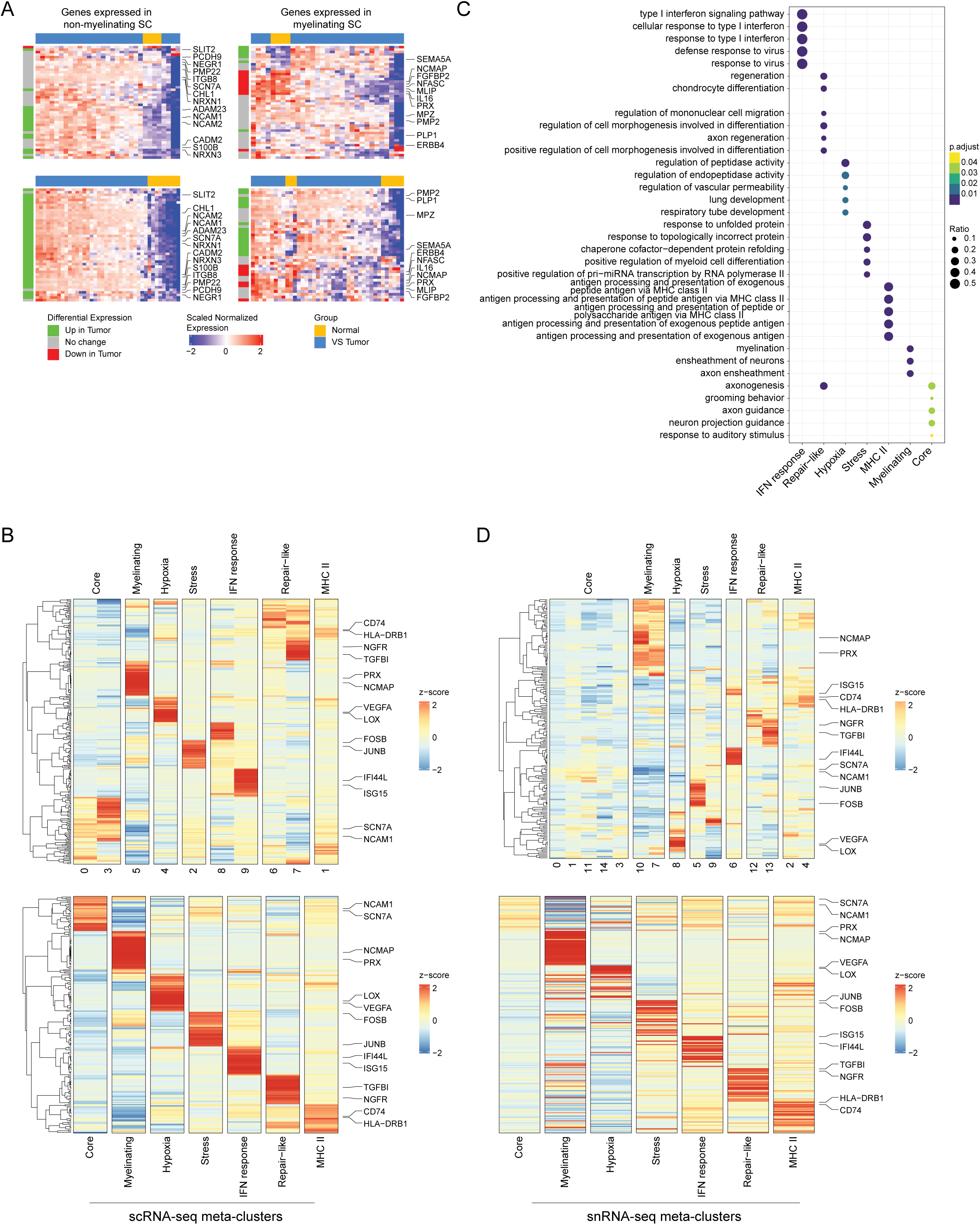
Supplementary VS-SC analysis. (A) Heatmaps comparing expression of top 50 differentially expressed genes (DEGs) in VS-nmSC and VS-myeSC to expression observed in microarray data of normal nerve and VS tumors (top, Zhao *et al.* (GSE108524); bottom, Torres-Martin *et al.* (GSE39645); see also Figure 2B). (B) Heatmap showing RPCA-based clustering results for VS SC subcluster (top) with hierarchical clustering of top 30 DEGs. Bottom heatmap shows final cluster-type labels and expression of cluster-defining genes. (C) Dot plot displaying results from GO BP enrichment analysis for top 25 DEGs from each VS-SC subtype. (D) Heatmaps showing RPCA-based clustering results for VS-SC from the snRNA-seq data, performed identically as in Figure S3B. There is strong correlation with gene expression and cluster assignment with samples from scRNA-seq analysis.

**Figure S4.**
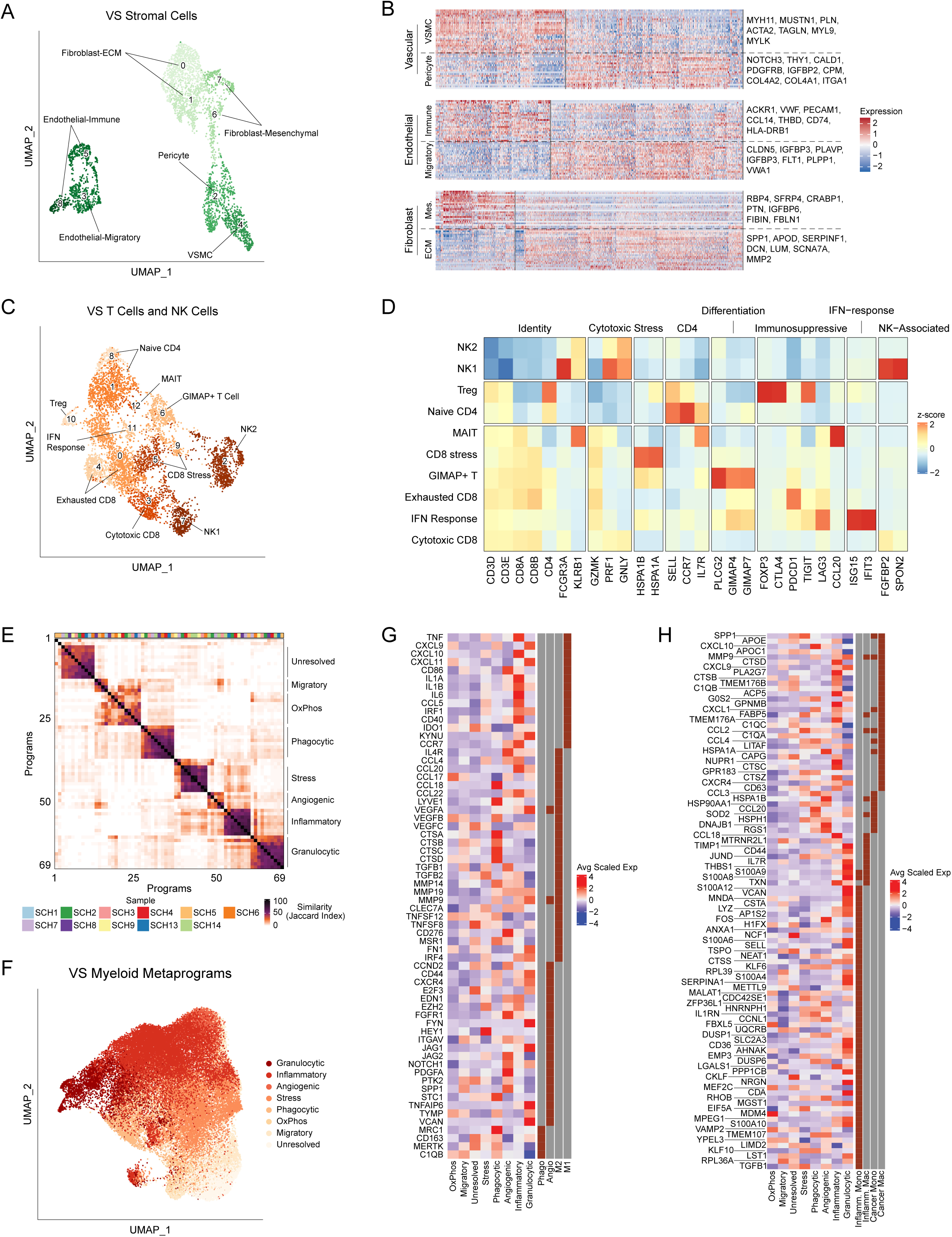
Classification of cell states in stromal and immune cell populations. (A) UMAP plot displaying stroma cell meta-cluster labels. (B) Heatmap showing representative gene expression for each stromal cell meta-cluster. (C) UMAP displaying NK and T cell meta-cluster labels. (D) Heatmap showing average expression across meta-clusters of known marker genes for NK and T cell phenotypes. (E) Heatmap displaying pairwise similarities between myeloid-cell programs, identified via NMF. Annotations on right designate meta-program (MP) labels. (F) Myeloid cells were scored for each MP identified in Figure S4A and assigned to the MP for which they scored highest. (G) Heatmap showing expression of cancer associated macrophage markers, as defined in a pan-cancer scRNA-seq analysis of tumor infiltrating macrophages^39^. (H) Heatmap showing expression of monocyte/macrophage markers expressed in the setting of inflammation and cancer, as defined in pan-tissue scRNA-seq analysis of myeloid cells^41^.

**Figure S5.**
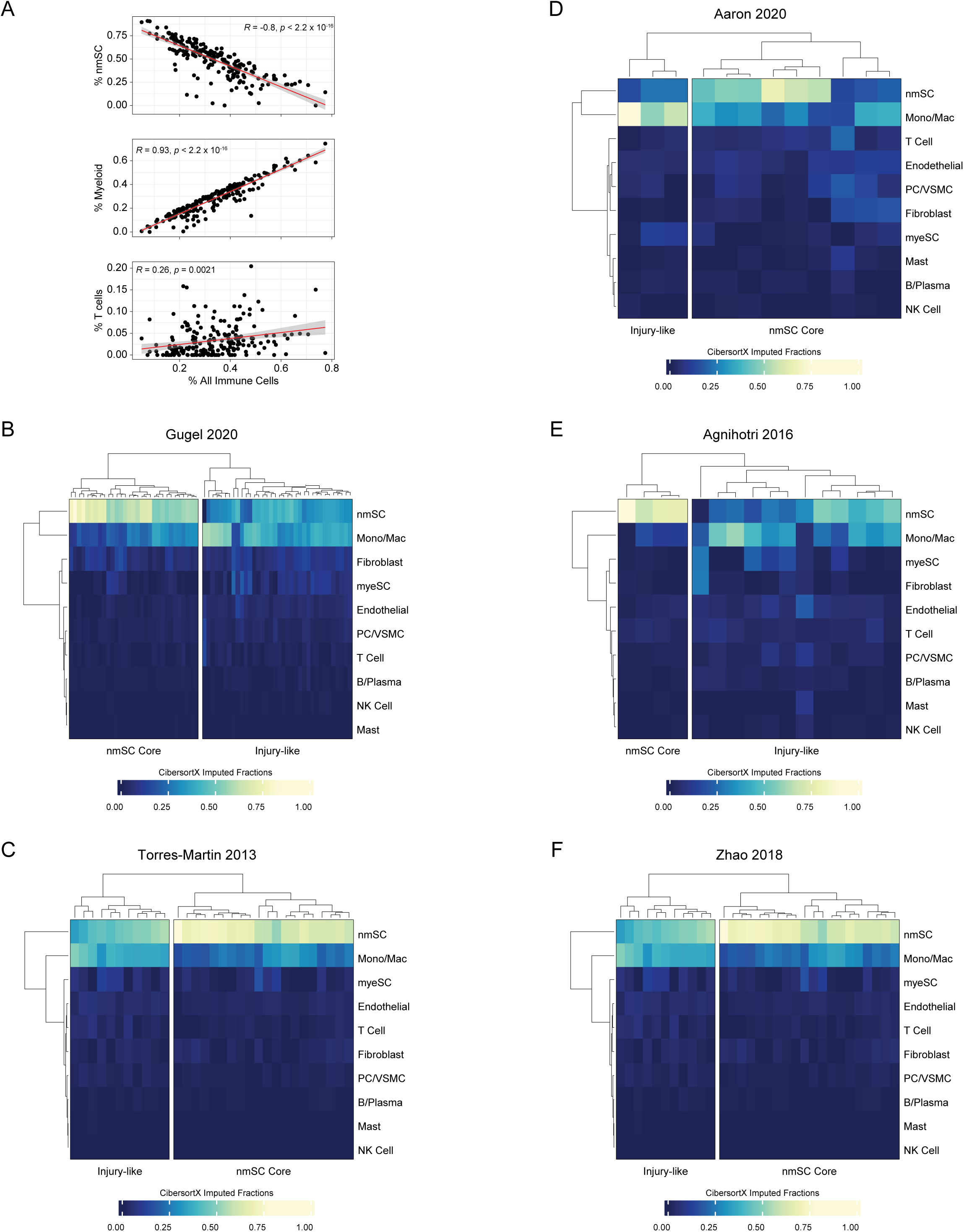
Classification of deconvolved bulk RNA expression data. (A) Correlation of imputed fractions of myeloid (top), T cell (middle), and nmSC (cells) with percentage of all immune cells in each deconvolved tumor sample. (B-F) Heatmaps displaying imputed cell fractions from CIBERSORTx deconvolution. VS tumors are classified into Injury-like and nmSC Core categories using hierarchical clustering of imputed cell fractions. BC, B cells; TC, T cells; NKC, Natural Killer Cells; PC/VSMC, pericyte/vascular smooth muscle cells; myeSC, myelinating Schwann cell; nmSC, non-myelinating Schwann cell.

**Figure S6.**
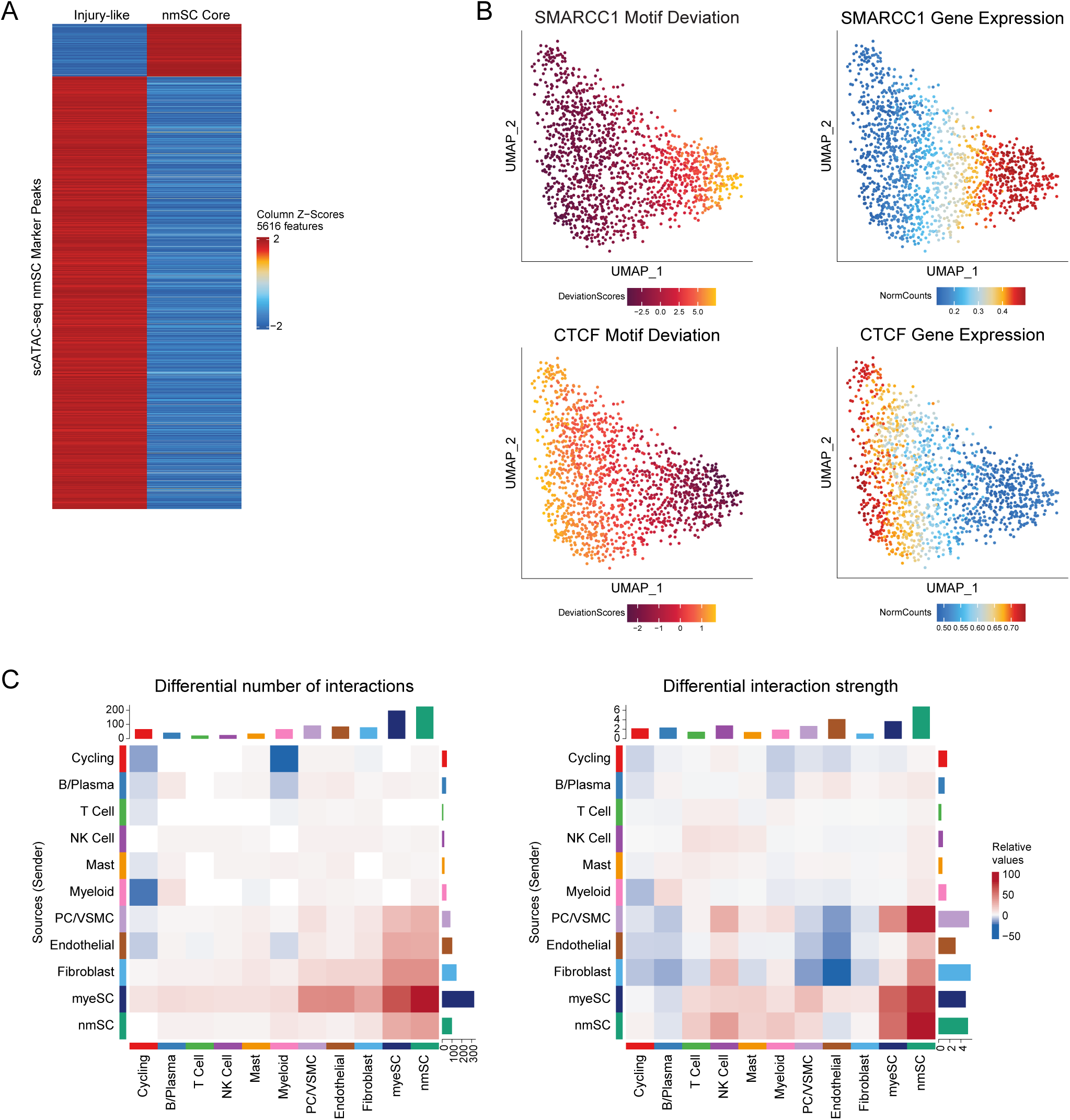
Supplemental scATAC-seq and ligand-receptor analyses. (A) Heatmap showing differentially accessible peaks (DAPs) identified 5616 statistically significant marker peaks with Log2FC ≥ 2 differentiating VS-SC in Injury-like and nmSC Core tumors. (B) UMAP of Motif Deviation and Gene Expression of select genes specific to nmSC Core and Injury-like VS SC. (C) Heatmap show differential number of interactions (left) and interaction strength (right), displayed as Injury-like VS relative to nmSC Core VS. The top colored bar plot represents the sum of column of values displayed (*i.e.*, incoming signals). The right colored bar plot represents the sum of row of values (*i.e.*, outgoing signals). Red and blue colors in the color scale represent increased and decreased signaling, respectively, in Injury-like VS nmSC Core tumors. BC, B cells; TC, T cells; NKC, Natural Killer Cells; PC/VSMC, pericyte/vascular smooth muscle cells; myeSC, myelinating Schwann cell; nmSC, non-myelinating Schwann cell.

## REFERENCES

1. Carlson, M. L. & Link, M. J. Vestibular Schwannomas. N Engl J Med 384, 1335–1348 (2021).

2. Starnoni, D. et al. Systematic review and meta-analysis of the technique of subtotal resection and stereotactic radiosurgery for large vestibular schwannomas: a “nerve-centered” approach. Neurosurgical Focus 44, E4 (2018).

3. Yang, I. et al. A comprehensive analysis of hearing preservation after radiosurgery for vestibular schwannoma: Clinical article. JNS 112, 851–859 (2010).

4. Coughlin, A. R., Willman, T. J. & Gubbels, S. P. Systematic Review of Hearing Preservation After Radiotherapy for Vestibular Schwannoma: Otology & Neurotology 39, 273–283 (2018).

5. Ahsan, S. F., Huq, F., Seidman, M. & Taylor, A. Long-term Hearing Preservation After Resection of Vestibular Schwannoma: A Systematic Review and Meta-analysis. Otology & Neurotology 38, 1505–1511 (2017).

6. Marinelli, J. P., Grossardt, B. R., Lohse, C. M. & Carlson, M. L. Prevalence of Sporadic Vestibular Schwannoma: Reconciling Temporal Bone, Radiologic, and Population-based Studies. Otology & Neurotology 40, 384–390 (2019).

7. Ahronowitz, I. et al. Mutational spectrum of the *NF2* gene: a meta-analysis of 12 years of research and diagnostic laboratory findings. Hum. Mutat. 28, 1–12 (2007).

8. Carlson, M. L. et al. Next Generation Sequencing of Sporadic Vestibular Schwannoma: Necessity of Biallelic NF2 Inactivation and Implications of Accessory Non-NF2 Variants. Otology & Neurotology 39, e860–e871 (2018).

9. Petrilli, A. M. & Fernández-Valle, C. Role of Merlin/NF2 inactivation in tumor biology. Oncogene 35, 537–548 (2016).

10. Neff, B. A. et al. Inhibition of MEK pathway in vestibular schwannoma cell culture. The Laryngoscope 122, 2269–2278 (2012).

11. Giovannini, M. et al. mTORC1 inhibition delays growth of neurofibromatosis type 2 schwannoma. Neuro-Oncology 16, 493–504 (2014).

12. Kaempchen, K. Upregulation of the Rac1/JNK signaling pathway in primary human schwannoma cells. Human Molecular Genetics 12, 1211–1221 (2003).

13. Blair, K. J. et al. EGF and bFGF Promote Invasion That Is Modulated by PI3/Akt Kinase and Erk in Vestibular Schwannoma: Otology & Neurotology 32, 308–314 (2011).

14. Zhou, L. et al. Merlin-Deficient Human Tumors Show Loss of Contact Inhibition and Activation of Wnt/β-Catenin Signaling Linked to the PDGFR/Src and Rac/PAK Pathways. Neoplasia 13, 1101-IN2 (2011).

15. Fuse, M. A. et al. Preclinical assessment of MEK1/2 inhibitors for neurofibromatosis type 2– associated schwannomas reveals differences in efficacy and drug resistance development. Neuro-Oncology 21, 486–497 (2019).

16. Goutagny, S. et al. Phase II study of mTORC1 inhibition by everolimus in neurofibromatosis type 2 patients with growing vestibular schwannomas. J Neurooncol 122, 313–320 (2015).

17. Karajannis, M. A. et al. Phase II study of everolimus in children and adults with neurofibromatosis type 2 and progressive vestibular schwannomas. Neuro-Oncology 16, 292–297 (2014).

18. Plotkin, S. R. et al. Multicenter, Prospective, Phase II and Biomarker Study of High-Dose Bevacizumab as Induction Therapy in Patients With Neurofibromatosis Type 2 and Progressive Vestibular Schwannoma. JCO 37, 3446–3454 (2019).

19. Qi, Z., Barrett, T., Parikh, A. S., Tirosh, I. & Puram, S. V. Single-cell sequencing and its applications in head and neck cancer. Oral Oncology 99, 104441 (2019).

20. Buenrostro, J. D. et al. Single-cell chromatin accessibility reveals principles of regulatory variation. Nature 523, 486–490 (2015).

21. Yim, A. K. Y. et al. Disentangling glial diversity in peripheral nerves at single-nuclei resolution. Nat Neurosci 25, 238–251 (2022).

22. Hung, G. et al. Immunohistochemistry study of human vestibular nerve schwannoma differentiation. Glia 38, 363–370 (2002).

23. Carr, M. J. et al. Mesenchymal Precursor Cells in Adult Nerves Contribute to Mammalian Tissue Repair and Regeneration. Cell Stem Cell 24, 240–256.e9 (2019).

24. Kalinski, A. L. et al. Analysis of the immune response to sciatic nerve injury identifies efferocytosis as a key mechanism of nerve debridement. eLife 9, e60223 (2020).

25. Wolbert, J. et al. Redefining the heterogeneity of peripheral nerve cells in health and autoimmunity. Proc. Natl. Acad. Sci. U.S.A. 117, 9466–9476 (2020).

26. Gerber, D. et al. Transcriptional profiling of mouse peripheral nerves to the single-cell level to build a sciatic nerve ATlas (SNAT). eLife 10, e58591 (2021).

27. Agnihotri, S. et al. The genomic landscape of schwannoma. Nat Genet 48, 1339–1348 (2016).

28. Müller, S., Cho, A., Liu, S. J., Lim, D. A. & Diaz, A. CONICS integrates scRNA-seq with DNA sequencing to map gene expression to tumor sub-clones. Bioinformatics 34, 3217–3219 (2018).

29. Gugel, I. et al. Contribution of mTOR and PTEN to Radioresistance in Sporadic and NF2-Associated Vestibular Schwannomas: A Microarray and Pathway Analysis. Cancers 12, 177 (2020).

30. Torres-Martin, M. et al. Microarray analysis of gene expression in vestibular schwannomas reveals SPP1/MET signaling pathway and androgen receptor deregulation. International Journal of Oncology 42, 848–862 (2013).

31. Zhao, Y. et al. Targeting the cMET pathway augments radiation response without adverse effect on hearing in NF2 schwannoma models. Proc Natl Acad Sci USA 115, E2077–E2084 (2018).

32. Helbing, D.-L., Schulz, A. & Morrison, H. Pathomechanisms in schwannoma development and progression. Oncogene 39, 5421–5429 (2020).

33. Hartlehnert, M. et al. Schwann cells promote post-traumatic nerve inflammation and neuropathic pain through MHC class II. Sci Rep 7, 12518 (2017).

34. Wang, Z. H., Walter, G. F. & Gerhard, L. The expression of nerve growth factor receptor on Schwann cells and the effect of these cells on the regeneration of axons in traumatically injured human spinal cord. Acta Neuropathol 91, 180–184 (1996).

35. Ding, D. et al. Runx2 was Correlated with Neurite Outgrowth and Schwann Cell Differentiation, Migration After Sciatic Nerve Crush. Neurochem Res 43, 2423–2434 (2018).

36. Wang, J.-B. et al. SPP1 promotes Schwann cell proliferation and survival through PKCα by binding with CD44 and αvβ3 after peripheral nerve injury. Cell Biosci 10, 98 (2020).

37. Curtis, R. et al. GAP-43 is expressed by nonmyelin-forming Schwann cells of the peripheral nervous system. Journal of Cell Biology 116, 1455–1464 (1992).

38. Hansen, M. R., Roehm, P. C., Chatterjee, P. & Green, S. H. Constitutive neuregulin-1/ErbB signaling contributes to human vestibular schwannoma proliferation. Glia 53, 593–600 (2006).

39. Cheng, S. et al. A pan-cancer single-cell transcriptional atlas of tumor infiltrating myeloid cells. Cell 184, 792–809.e23 (2021).

40. Kinker, G. S. et al. Pan-cancer single-cell RNA-seq identifies recurring programs of cellular heterogeneity. Nat Genet 52, 1208–1218 (2020).

41. Mulder, K. et al. Cross-tissue single-cell landscape of human monocytes and macrophages in health and disease. Immunity 54, 1883–1900.e5 (2021).

42. Ginhoux, F., Schultze, J. L., Murray, P. J., Ochando, J. & Biswas, S. K. New insights into the multidimensional concept of macrophage ontogeny, activation and function. Nat Immunol 17, 34–40 (2016).

43. Newman, A. M. et al. Determining cell type abundance and expression from bulk tissues with digital cytometry. Nat Biotechnol 37, 773–782 (2019).

44. Aaron, K. A. et al. What Genes Can Tell: A Closer Look at Vestibular Schwannoma. Otology & Neurotology 41, 522–529 (2020).

45. Arthur-Farraj, P. J. et al. Changes in the Coding and Non-coding Transcriptome and DNA Methylome that Define the Schwann Cell Repair Phenotype after Nerve Injury. Cell Reports 20, 2719–2734 (2017).

46. Li, M., Banton, M. C., Min, Q., Parkinson, D. B. & Dun, X. Meta-Analysis Reveals Transcription Factor Upregulation in Cells of Injured Mouse Sciatic Nerve. Front. Cell. Neurosci. 15, 688243 (2021).

47. Wang, J. et al. CTCF-mediated chromatin looping in EGR2 regulation and SUZ12 recruitment critical for peripheral myelination and repair. Nat Commun 11, 4133 (2020).

48. Ma, K. H., Hung, H. A. & Svaren, J. Epigenomic Regulation of Schwann Cell Reprogramming in Peripheral Nerve Injury. Journal of Neuroscience 36, 9135–9147 (2016).

49. Jin, S. et al. Inference and analysis of cell-cell communication using CellChat. Nat Commun 12, 1088 (2021).

50. Pruenster, M. et al. The Duffy antigen receptor for chemokines transports chemokines and supports their promigratory activity. Nat Immunol 10, 101–108 (2009).

51. Ko, K. R., Lee, J., Lee, D., Nho, B. & Kim, S. Hepatocyte Growth Factor (HGF) Promotes Peripheral Nerve Regeneration by Activating Repair Schwann Cells. Sci Rep 8, 8316 (2018).

52. de Vries, W. M., Briaire-de Bruijn, I. H., van Benthem, P. P. G., van der Mey, A. G. L. & Hogendoorn, P. C. W. M-CSF and IL-34 expression as indicators for growth in sporadic vestibular schwannoma. Virchows Arch 474, 375–381 (2019).

53. Trias, E. et al. Schwann cells orchestrate peripheral nerve inflammation through the expression of CSF1, IL-34, and SCF in amyotrophic lateral sclerosis. Glia 68, 1165–1181 (2020).

54. Martini, R., Fischer, S., López-Vales, R. & David, S. Interactions between Schwann cells and macrophages in injury and inherited demyelinating disease. Glia 56, 1566–1577 (2008).

55. Prueter, J., Norvell, D. & Backous, D. Ki-67 index as a predictor of vestibular schwannoma regrowth or recurrence. J. Laryngol. Otol. 133, 205–207 (2019).

56. Lewis, D. et al. Inflammation and vascular permeability correlate with growth in sporadic vestibular schwannoma. Neuro-Oncology 21, 314–325 (2019).

57. Breun, M. et al. CXCR4: A new player in vestibular schwannoma pathogenesis. Oncotarget 9, (2018).

58. Hannan, C. J. et al. The inflammatory microenvironment in vestibular schwannoma. Neuro-Oncology Advances 2, vdaa023 (2020).

59. de Vries, M. et al. Tumor-Associated Macrophages Are Related to Volumetric Growth of Vestibular Schwannomas: Otology & Neurotology 34, 347–352 (2013).

60. Kandathil, C. K., Cunnane, M. E., McKenna, M. J., Curtin, H. D. & Stankovic, K. M. Correlation Between Aspirin Intake and Reduced Growth of Human Vestibular Schwannoma: Volumetric Analysis. Otology & Neurotology 37, 1428–1434 (2016).

61. Hagan, N. et al. CSF1R signaling is a regulator of pathogenesis in progressive MS. Cell Death Dis 11, 904 (2020).

62. Fleming, S. J., et al. Unsupervised removal of systematic background noise from droplet-based single-cell experiments using CellBender. http://biorxiv.org/lookup/doi/10.1101/791699 (2019) doi:10.1101/791699.

63. Wolock, S. L., Lopez, R. & Klein, A. M. Scrublet: Computational Identification of Cell Doublets in Single-Cell Transcriptomic Data. Cell Systems 8, 281–291.e9 (2019).

64. Wolf, F. A., Angerer, P. & Theis, F. J. SCANPY: large-scale single-cell gene expression data analysis. Genome Biol 19, 15 (2018).

65. Satija, R., Farrell, J. A., Gennert, D., Schier, A. F. & Regev, A. Spatial reconstruction of single-cell gene expression data. Nat Biotechnol 33, 495–502 (2015).

66. Granja, J. M. et al. ArchR is a scalable software package for integrative single-cell chromatin accessibility analysis. Nat Genet 53, 403–411 (2021).

67. Hao, Y. et al. Integrated analysis of multimodal single-cell data. Cell 184, 3573–3587.e29 (2021).

68. Beaubier, N. et al. Clinical validation of the tempus xT next-generation targeted oncology sequencing assay. Oncotarget 10, 2384–2396 (2019).

69. Browaeys, R., Saelens, W. & Saeys, Y. NicheNet: modeling intercellular communication by linking ligands to target genes. Nat Methods 17, 159–162 (2020).

