## Supplementary Tables for "Single-cell multi-omic analysis of the vestibular schwannoma ecosystem uncovers a nerve injury-like state"

**Supplementary Table 1. scRNA-seq Patient Characteristics**

| Subject | Technique | scATAC | Sex | Age | Size | Consistenc | Subjective_ | Previous_T | Dysequilibi | Trigem | WRS | PT500 | PT1000 | PT2000 | PTA | AAO-HNS | Resection | Adherent | FN_Stim | Postop_FN | Postop_Recurrence |
|---|---|---|---|---|---|---|---|---|---|---|---|---|---|---|---|---|---|---|---|---|---|
| SCH1 | scRNA | No | M | 62 | 2.6 | Solid | Yes | No | Yes | No | 96 | 10 | 20 | 30 | 20 | A | GTR | No | Yes | Yes | No |
| SCH2 | scRNA | No | F | 88 | 3.4 | Cystic | Yes | No | Yes | Yes | 40 | 100 | 100 | 100 | 100 | D | STR | Yes | Yes | No | No |
| SCH3 | scRNA | No | F | 44 | 2.5 | Cystic | Yes | No | No | No | 0 | 100 | 100 | 100 | 100 | D | STR | Yes | Yes | No | No |
| SCH4 | scRNA | Yes | F | 59 | 3.7 | Solid | Yes | No | No | Yes | 0 | 75 | 80 | 75 | 76.66667 | D | STR | Yes | Yes | Yes | Yes |
| SCH5 | scRNA | Yes | M | 65 | 2 | Solid | Yes | No | No | No | 4 | 60 | 55 | 85 | 70 | D | GTR | No | Yes | No | No |
| SCH6 | scRNA | Yes | M | 43 | 3.1 | Solid | No | No | Yes | No | 96 | 20 | 20 | 25 | 21.66667 | A | GTR | No | Yes | Yes | No |
| SCH7 | scRNA | Yes | F | 51 | 1.3 | Solid | Yes | No | Yes | No | 40 | 20 | 65 | 60 | 48.33333 | D | GTR | No | Yes | No | No |
| SCH8 | scRNA | No | F | 64 | 1.4 | Solid | Prior Tx | Yes | No | No | 0 | 100 | 100 | 100 | 100 | D | GTR | No | Yes | No | No |
| SCH9 | scRNA | Yes | M | 72 | 0.61 | Solid | Yes | No | Yes | No | 0 | 100 | 100 | 100 | 100 | D | GTR | No | Yes | No | No |
| SCH13 | scRNA | Yes | F | 67 | 3.6 | Cystic | Prior Tx | Yes | No | No | 0 | 100 | 100 | 100 | 100 | D | STR | Yes | Yes | No | No |
| SCH14 | scRNA | Yes | F | 68 | 2.1 | Solid | Prior Tx | Yes | No | Yes | 0 | 100 | 100 | 100 | 100 | D | STR | Yes | Yes | Yes | No |
| SCH18 | snRNA | No | M | 66 | 1.9 | Solid | Prior Tx | No | No | No | 0 | 100 | 100 | 100 | 100 | D | GTR | Yes | Yes | No | No |
| SCH20 | snRNA | No | F | 36 | 2.1 | Cystic | Yes | No | No | No | 82 | 20 | 15 | 35 | 23.33333 | A | STR | Yes | Yes | No | No |
| SCH21 | snRNA | No | M | 34 | 3.2 | Cystic | Yes | No | Yes | Yes | 0 | 100 | 100 | 100 | 100 | D | STR | Yes | Yes | Yes | No |
| SCH22 | snRNA | No | M | 37 | 3.9 | Solid | Yes | No | No | No | 45 | 100 | 100 | 100 | 100 | D | STR | Yes | Yes | No | No |

**Supplementary Table 2. Bulk RNA-seq patient characteristics**

| ID | Sex | Age | Side | HL | Tinnitus | Vertigo | Size | NF2 | Recurrence | Prior_Sx | Prior_SRS | Postop_SRS | EOR | Postop_Recur |
|---|---|---|---|---|---|---|---|---|---|---|---|---|---|---|
| 15-04-001 | M | 24 | Right | Y | N | N | 3.17 | N | Y | Y | N | N | GTR | N |
| 17-04-006 | M | 18 | Left | Y | Y | N | 4.11 | Y | N | N | Y | N | STR | N |
| 17-04-007 | F | 37 | Left | Y | Y | N | 1.55 | N | N | N | N | N | GTR | N |
| 19-04-001 | F | 40 | Left | N | Y | N | 2.45 | N | N | N | N | N | GTR | N |
| 19-04-002 | F | 45 | Left | Y | N | N | 2.59 | N | Y | Y | N | Y | STR | Y |
| 19-04-003 | M | 67 | Right | Y | N | N | 3.74 | N | N | N | N | N | GTR | N |
| 19-04-004 | F | 19 | Left | Y | N | N | 3.09 | N | N | N | N | N | STR | N |
| 19-04-005 | F | 48 | Right | Y | Y | Y | 2.87 | N | N | N | N | N | GTR | N |
| 19-04-007 | F | 45 | Left | Y | N | N | 1.82 | N | N | N | N | N | STR | N |
| 19-04-008 | F | 45 | Right | Y | Y | Y | 1.25 | N | N | N | N | N | GTR | N |
| 19-04-009 | M | 33 | Right | Y | N | Y | 4.62 | N | N | N | N | N | STR | Y |
| 19-04-010 | F | 32 | Left | Y | Y | N | 1.98 | N | N | N | N | N | GTR | N |
| 20-04-001 | F | 42 | Right | Y | N | N | 3.41 | N | N | N | N | N | GTR | N |
| 20-04-002 | M | 49 | Left | Y | Y | Y | 1.44 | N | N | N | N | N | GTR | N |
| 20-04-005 | M | 38 | Right | Y | Y | Y | 1.63 | N | N | N | N | N | GTR | N |
| 20-04-008 | F | 52 | Left | Y | Y | N | 1.23 | N | N | N | N | N | GTR | N |
| 21-04-001 | F | 65 | Left | Y | Y | N | 2.68 | N | N | N | N | N | GTR | N |
| 21-04-002 | F | 55 | Left | Y | Y | N | 1.23 | N | N | N | N | N | GTR | N |
| 21-04-003 | F | 22 | Right | Y | Y | N | 2.96 | N | N | N | N | N | GTR | N |
| 21-04-004 | F | 45 | Left | Y | N | Y | 1.63 | N | N | N | N | N | GTR | N |
| 21-04-005 | F | 60 | Right | Y | N | N | 2.47 | N | N | N | N | N | STR | N |
| 21-04-006 | M | 26 | Left | N | Y | Y | 2.85 | N | N | N | N | N | GTR | N |

**Supplementary Table 3. Sequencing QC Data**

| Sample | Technique | Estimated Nu | Mean Reads p | Median Genes | Number of Re | Valid Barcode | Sequencing Sa | Q30 Bases in B | Q30 Bases in R | Q30 Bases in U | Reads Mappe | Reads Mappe | Reads Mappe | Reads Mappe | Reads Mappe | Reads Mappe | Reads Mappe | Fraction Read | Total Genes D | Median UMI Counts per Cell |
|---|---|---|---|---|---|---|---|---|---|---|---|---|---|---|---|---|---|---|---|---|
| SCH1 | scRNA | 8,095 | 74,232 | 1,889 | 600,914,469 | 97.00% | 76.80% | 93.80% | 92.30% | 93.50% | 96.90% | 93.30% | 10.00% | 27.30% | 55.90% | 52.80% | 1.80% | 88.10% | 27,639 | 4,461 |
| SCH2 | scRNA | 8,334 | 58,843 | 2,407 | 490,399,563 | 95.80% | 65.60% | 94.50% | 90.80% | 94.00% | 94.70% | 91.40% | 7.50% | 30.60% | 53.40% | 50.00% | 2.10% | 92.20% | 26,576 | 8,383 |
| SCH3 | scRNA | 2,384 | 218,762 | 2,570 | 521,528,898 | 96.70% | 88.50% | 94.50% | 90.10% | 94.10% | 94.10% | 90.90% | 8.30% | 36.50% | 46.20% | 42.60% | 2.20% | 95.30% | 25,739 | 8,101 |
| SCH4 | scRNA | 29,098 | 20,875 | 1,359 | 607,421,636 | 89.30% | 43.70% | 94.90% | 93.60% | 95.10% | 95.70% | 90.60% | 8.40% | 30.60% | 51.60% | 46.80% | 3.30% | 80.70% | 28,904 | 3,271 |
| SCH5 | scRNA | 5,645 | 65,365 | 2,010 | 368,987,075 | 95.90% | 77.80% | 96.00% | 93.90% | 95.80% | 97.40% | 94.50% | 6.60% | 25.70% | 62.20% | 57.80% | 2.90% | 90.00% | 25,411 | 6,173 |
| SCH6 | scRNA | 10,719 | 33,232 | 1,774 | 356,216,174 | 96.80% | 62.30% | 95.90% | 93.70% | 95.80% | 96.90% | 91.60% | 7.70% | 27.80% | 56.10% | 52.50% | 2.20% | 84.20% | 26,783 | 4,196 |
| SCH7 | scRNA | 5,411 | 60,702 | 2,500 | 328,461,525 | 97.40% | 63.50% | 95.90% | 93.30% | 95.80% | 97.30% | 90.10% | 5.60% | 22.50% | 61.90% | 58.20% | 2.30% | 70.50% | 25,460 | 8,240 |
| SCH8 | scRNA | 7,162 | 60,367 | 2,872 | 432,353,933 | 98.00% | 63.00% | 95.60% | 92.60% | 95.20% | 97.30% | 94.60% | 6.20% | 21.10% | 67.30% | 64.30% | 1.40% | 89.60% | 27,097 | 11,312 |
| SCH9 | scRNA | 6,529 | 63,331 | 1,377 | 413,491,191 | 97.90% | 81.50% | 95.50% | 93.50% | 95.10% | 96.30% | 86.90% | 9.50% | 25.40% | 52.10% | 48.60% | 1.80% | 81.70% | 26,177 | 3,248 |
| SCH13 | scRNA | 15,743 | 28,223 | 2,147 | 444,317,084 | 97.50% | 43.10% | 94.90% | 91.80% | 94.80% | 96.40% | 94.40% | 6.30% | 30.60% | 57.50% | 53.60% | 2.00% | 92.50% | 26,474 | 6,894 |
| SCH14 | scRNA | 7,842 | 63,169 | 3,095 | 495,374,001 | 98.10% | 56.30% | 92.70% | 89.60% | 92.50% | 95.90% | 93.80% | 8.80% | 30.20% | 54.80% | 51.40% | 1.50% | 93.80% | 25,658 | 12,459 |
| SCH18 | snRNA | 19,791 | 15,714 | 1,682 | 310,994,267 | 95.00% | 37.80% | 95.00% | 89.10% | 94.90% | 94.20% | 90.90% | 5.70% | 56.70% | 28.50% | 47.10% | 37.60% | 85.50% | 32,583 | 2,593 |
| SCH20 | snRNA | 15,782 | 22,190 | 1,495 | 350,203,824 | 95.20% | 59.50% | 94.90% | 89.80% | 94.80% | 94.40% | 91.50% | 5.60% | 61.40% | 24.50% | 45.00% | 40.40% | 81.40% | 31,273 | 2,383 |
| SCH21 | snRNA | 19,353 | 17,990 | 1,830 | 348,169,608 | 95.80% | 44.20% | 94.90% | 89.70% | 94.80% | 92.00% | 88.10% | 4.70% | 54.70% | 28.70% | 52.20% | 30.60% | 84.60% | 31,514 | 3,099 |
| SCH22 | snRNA | 18,868 | 24,051 | 2,000 | 453,800,288 | 95.50% | 53.90% | 94.30% | 88.60% | 93.90% | 92.60% | 89.20% | 5.70% | 54.80% | 28.70% | 54.30% | 28.60% | 82.70% | 31,457 | 3,399 |

**Supplementary Table 4. CellBender Parameters**

| Sample | Expected Cells | Total Droplets | Low Count Th | FPR | Epochs | Learning Rate | Z-dims | Z-layers | Model | Training Fraction |
|---|---|---|---|---|---|---|---|---|---|---|
| SCH1 | 7000 | 800000 | 25 | 0.01 | 150 | 0.000001 | 100 | 500 | Full | 0.9 |
| SCH2 | 8000 | 800000 | 15 | 0.01 | 150 | 0.000002 | 100 | 500 | Full | 0.9 |
| SCH3 | 2500 | 800000 | 10 | 0.01 | 150 | 0.000001 | 200 | 1000 | Full | 0.9 |
| SCH4 | 5000 | 800000 | 25 | 0.01 | 150 | 0.000001 | 100 | 500 | Full | 0.9 |
| SCH5 | 5000 | 800000 | 15 | 0.01 | 150 | 0.000001 | 100 | 500 | Full | 0.9 |
| SCH6 | 7000 | 800000 | 10 | 0.01 | 150 | 0.000001 | 200 | 500 | Full | 0.9 |
| SCH7 | 4000 | 800000 | 10 | 0.01 | 150 | 0.000002 | 100 | 500 | Full | 0.9 |
| SCH8 | 8000 | 600000 | 10 | 0.01 | 150 | 0.000001 | 100 | 500 | Full | 0.9 |
| SCH9 | 5000 | 700000 | 5 | 0.01 | 150 | 0.0000005 | 150 | 1000 | Full | 0.9 |
| SCH13 | 10000 | 900000 | 10 | 0.01 | 150 | 0.000001 | 100 | 500 | Full | 0.9 |
| SCH14 | 9000 | 500000 | 5 | 0.01 | 150 | 0.000002 | 100 | 500 | Full | 0.9 |
| SCH20 | 18000 | 750000 | 10 | 0.01 | 150 | 0.000001 | 100 | 500 | full | 0.9 |
| SCH21 | 18000 | 900000 | 10 | 0.01 | 150 | 0.0000005 | 100 | 500 | full | 0.9 |
| SCH22 | 20000 | 900000 | 10 | 0.01 | 150 | 0.0000005 | 100 | 500 | full | 0.9 |
